# Viral evolution of T1L mammalian orthoreovirus enhances breast cancer stem-like cell killing

**DOI:** 10.64898/2025.12.10.693591

**Authors:** Natasha I. Roman Ortiz, Ella Boytim, Kylie Miller, Justin Hwang, Pranav Danthi, Julie H Ostrander

## Abstract

**Background:** Breast cancer is the second leading cause of death in women in the US. Among the different subtypes of breast cancer, estrogen receptor positive (ER+) has a more favorable prognosis; however, patients can experience cancer recurrence 10 years or more after initial diagnosis. Breast cancer stem-like cells (BCSCs) are slowly proliferating cells that drive metastasis and resistance to therapies that target rapidly proliferating tumor cells. BCSC self-renewal and survival pathways make them difficult to eliminate; no selective therapeutics currently exist to target them. Our research aims to use mammalian orthoreovirus (MRV) as an oncolytic agent to eliminate BCSCs.

**Methods:** We evaluated the effects of three different MRV strains (T3D, T1L and R2) under both 2D and 3D culture conditions (which enrich for BCSCs) in parental and paclitaxel resistant (TaxR) MCF7 cells using cell viability assays, flow cytometry, western blotting, and tumorsphere assays. We identified differentially expressed genes in response to T1L by RNA sequencing. Novel MRV strains were developed by serial passaging of the parental T1L strain in TaxR BCSCs, leading to the isolation and sequencing of three distinct MRV clones. The novel MRV clones were assessed for their oncolytic effects using the aforementioned techniques alongside inhibitor-based studies targeting cell death pathways.

**Results:** Notably, at least two clones (T1L SP B and T1L SP C) demonstrated enhanced oncolytic potency compared to the parent T1L strain against MCF7 TaxR BCSCs, even though their infectivity and viral protein synthesis was reduced. Whole genome sequencing of the viral clones identified changes in viral protein sequences that correlate with enhanced potency against BCSCs. T1L, T1L SP B and T1L SP C induced apoptosis; however, T1L SP B may induce a more immunogenic form of cell death in the TaxR cells, as indicated by the increased release of extracellular ATP (eATP), a damage-associated molecular pattern (DAMP), which is associated with immunogenic cell death.

**Conclusions:** In summary, we developed novel oncolytic viruses promote oncolysis in therapy-resistant BCSCs. We found that T1L SP B induces apoptosis in a more immunogenic manner, highlighting the potential of these viruses to eliminate BCSCs and stimulate the immune system.

## Background

Estrogen receptor positive (ER+) breast cancer is the most prevalent subtype of breast cancer, affecting two-thirds of patients, and is characterized by a prolonged risk of recurrence. For this subtype of breast cancer, recurrence can occur 10 years or more after initial diagnosis (1). Patients with advanced ER+ breast cancer are treated with a combination of CDK4/6 inhibitors (e.g. palbociclib, abemaciclib), and aromatase inhibitors (e.g. anastrazole, letrozole) or selective estrogen degraders (e.g. fulvestrant) (1). While these therapies prolong the lives of women with advanced, metastatic ER+ breast cancer, no curative treatments are available.

Accumulating evidence has shown that breast cancer stem-like cells (BCSCs) drive tumor initiation, progression and metastasis (2,3). BCSCs represent a minority subpopulation of cells that are drug resistant and are characterized by increased stemness markers and enhanced tumorigenic potential, which may contribute to breast cancer recurrence or relapse following standard of care therapy (3). These cells can self-renew and regenerate the cancer cell population at distant parts of the body (i.e. metastasis). BCSCs are typically characterized by expression of CD44^hi^/CD24^lo^ markers, elevated aldehyde dehydrogenase (ALDH1) activity, activation of survival pathways (such as Notch, Hedgehog, Hippo, and Wnt), and high expression of anti-apoptotic proteins including Survivin and Bcl-xL (2,4,5). To date, there are no FDA-approved treatments that selectively target BCSCs. BCSC drug resistance can result from the activity of transporters that efflux therapeutic agents out of the cell, elevated levels of ALDH1 which detoxifies anti-cancer substances, enhanced DNA damage repair activity, and high expression of stemness genes and anti-apoptotic proteins (2). A major challenge in targeting BCSCs is their biological heterogeneity and plasticity (6). In addition, BCSCs share similar signaling pathways with normal stem cells (7). Universal, reliable biomarkers for BCSCs have not been identified, which complicates efforts in the development of BCSC-targeted treatments.

Oncolytic viruses (OV) are becoming a promising therapeutic approach to target cancer cells, with the FDA approving Talimogene laherparepvec (T-VEC or Imlygic), a genetically modified herpes simplex virus, for the treatment of advanced melanoma in 2015 (8). OVs directly kill tumor cells while sparing normal cells (9,10). OVs can directly kill cancer cells due to their favorable environment, such as defects in tumor-suppressor genes, reduced activity of antiviral pathways in tumor cells, interactions with specific receptors, or the engineering of viral vectors with targeted gene deletions (10). Genetic engineering also allows OVs to carry exogenous genes that enhance tumor targeting, replication, and lysis, while also stimulating antigen-specific immune responses, highlighting their potential as immunotherapy agents (9,10).

Mammalian orthoreovirus (MRV), or reovirus, is an icosahedral, non-enveloped double stranded RNA (dsRNA) OV composed of ten genome segments (three large, three medium, four small) (9). The ten genome segments encode for eight structural proteins (σ1, σ2, σ3, μ1, μ2, λ1, λ2, and λ3) and four nonstructural proteins (σ1s, σNS, μNS and μNSC) (11). MRV can infect most mammals; humans are typically infected in early childhood without significant adverse pathology (12). MRV preferentially replicates in transformed cells while sparing normal cells (13). Four distinct serotypes have been identified: Type 1 Lang (T1L), Type 2 Jones (T2J), Type 3 strains (including Abney T3A and Dearing strains T3D), and Type 4 Ndelle (14–16). While all strains express the same set of gene products, there are polymorphic differences between them (17,18). Among these prototypes, T3D is the most studied and has been evaluated in clinical trials for its oncolytic properties, while T2J is the least characterized. To initiate infection, reovirus particles attach to a variety of receptors, such as Junctional Adhesion Molecule A (JAM-A), sialic acid, PirB, Nrp1, NgR1, and integrins (14,19–22). The particles are endocytosed, and the outer protein shell is disassembled through the action of acid-activated host proteases (23). Conformational changes in the remaining outer capsid proteins allow the particle to bypass the endosomal membrane (24–26). Once the uncoated particles reach the cytoplasm, the viral core becomes transcriptionally active, allowing virus replication to commence (24–26).

MRV (Reolysin, strain T3D) has been investigated in Phase I-III clinical trials in different cancer types (head and neck, pancreatic, prostate, breast, and others). Results in clinical trials have shown that MRV is safe, and the primary side effects are low-grade, flu-like symptoms (27). MRV combined with chemotherapy is well tolerated and may have synergistic benefits (28–32). In 2015, the FDA approved Reolysin as an orphan drug for the treatment for ovarian, pancreatic, and other cancer types (33). In 2017, it received FDA fast-track designation for treating metastatic breast cancer (34). Additionally, Reolysin was shown to enhance anti-tumor immunity in high-grade gliomas and metastatic brain tumors (35).

Even though MRV has been shown to induce oncolysis in breast cancer cells, little is known about MRV’s effect in BCSCs. Marcato et al. demonstrated that MRV effectively induces tumor regression and targets BCSCs in human breast cancer xenografts in mice, with both BCSCs and non-BCSCs undergoing apoptosis (36). The findings suggest that MRV therapy could help reduce tumor cells and BCSCs in breast cancer. A previous study isolated a novel MRV strain (R2), through serial passaging of three different parent MRV strains (T3D, T1L and T2J) in MDA-MB-231 triple-negative breast cancer (TNBC) cells. R2 was found to contain genome segments from T3D and T1L along with several point mutations, and it showed improved infection and killing in TNBC cells (37), but the effect of R2 on BCSCs was not tested. In this study, we compared the effects of three different MRV strains (T3D, T1L, R2) in ER+ breast cancer cell lines. We tested T3D, T1L, and R2 in standard cell cultures and BCSC-enriched tumorspheres and found that they were generally less effective against BCSCs. To improve BCSC killing, we performed serial passaging of T1L in BCSC-enriched conditions and isolated three novel viral strains. Two strains showed stronger killing of therapy-resistant BCSCs than the parent MRV. Our approach demonstrates a powerful tool to improve MRV infectivity and cytotoxicity for the advancement of oncolytic therapy in treating advanced ER+ breast cancer.

## Methods

### Cell lines

The human breast cancer cell line MCF7 was obtained from ATCC (Manassas, VA) and the MCF7 paclitaxel resistant (TaxR) cell line was established as described in earlier studies (38,39). The normal mammary epithelial cell line MCF710A was obtained from ATCC. UCD4 and UCD12 cell lines were obtained from Dr. Carol Sartorius (University of Colorado Anschutz Medical Campus). UCD4 TamR (tamoxifen resistant) and UCD12 PalboR (palbociclib resistant) were obtained by continuous culturing of UCD4 and UCD12 in increasing concentrations of tamoxifen or palbociclib over a 6-9 month period. The L929 cell line was provided by Dr. Pranav Danthi (Indiana University, Bloomington). Both MCF7 and TaxR cells were grown in MEM-alpha media (Thermo Fisher Scientific, Waltham, MA). MEM-alpha media was supplemented with 5% FBS (Cytiva-HyClone, Logan, UT), 12 mM HEPES (Thermo Fisher Scientific, Waltham, MA), 1mM Sodium Pyruvate (Thermo Fisher Scientific, Waltham, MA), 1X MEM Non-Essential Amino Acids (Thermo Fisher Scientific, Waltham, MA), 1 µg/mL insulin (Thermo Fisher Scientific, Waltham, MA), 1 µg/mL hydrocortisone (Millipore Sigma, Burlington, MA), and 12.5 ng/mL EGF (Millipore Sigma, Burlington, MA). MCF7 TaxR cells were cultured with 2 μM paclitaxel (Invitrogen, Waltham, MA). MCF10A cells were grown in DMEM-F12 (Corning, Incorporated, Corning, NY) media supplemented with 5% horse serum (Corning, Incorporated, Corning, NY), 10 μg/mL insulin, 20 ng/mL EGF, 100 ng/mL cholera toxin (Millipore Sigma, Burlington, MA), and 0.47 µg/mL hydrocortisone. The UCD cells were cultured in DMEM-F12, 10% FBS, 100 ng/mL cholera toxin and 1 nM insulin. UCD4 TamR cells were cultured with 1 μM tamoxifen (Millipore Sigma, Burlington, MA) and UCD4 PalboR were cultured with 1 μM palbociclib (Fisher Scientific, Waltham, MA). L929 cells were cultured with MEM media (Corning, Incorporated, Corning, NY) supplemented with 5% FBS and 2 mM glutamine (Thermo Fisher Scientific, Waltham, MA). All cell lines were authenticated prior to the initiation of experiments. Authenticated cell lines were expanded and frozen down in multiple aliquots. Experiments were performed within 20 passages from thaw. All cell lines were tested for mycoplasma every 6 months.

### Cell viability Assay

Cells were plated at 2,000-7,000 cells per well in 96-well plates (Corning, Incorporated, Corning, NY) with 100 µL of complete media (appropriate for the specific cell line). After 24h, cells were infected with MRV at 100 PFU/cell. Media was removed and MRV was diluted in serum-free DMEM 1X (Corning, Incorporated, Corning, NY) for 2D conditions, to a final volume of 25 µL. MRV was added to the corresponding wells and incubated for 1h at room temperature (RT) with rocking. Complete media was added (100 µL) and cells were incubated at 37°C for 4 days. After 4 days of MRV infection, 10% (in total media) of alamarBlue (Bio-Rad, Hercules, CA) cell viability reagent was added to each well and incubated at 37°C for 2h. Fluorescence was measured at 540/620 nm using the BioTek Synergy 2 Microplate Reader and Gen5 software (BioTek Instruments, Inc, Winooski, VT). Data was analyzed, normalizing to mock control, to obtain the % decrease in cell viability.

For inhibitor studies, cells were seeded and infected as described above, but using a final volume of 12.5 µL. Twenty-four hours post infection, Q-VD-Oph (20 µM) (SelleckChem, Houston, TX) or necrostatin-1 (50 µM) (Tocris Bioscience Fisher Scientific, Waltham, MA) were added at 12.5 µL per well, followed by incubation at 37°C for 48h, and then retreated with the same inhibitor concentrations, cell viability was assessed with the alamarBlue assay the following day.

### Tumorsphere Assays

After trypsinization, single cell suspensions were obtained by passing cells through a 40-µM cell strainer (Corning, Incorporated, Corning, NY). Cells were plated at 1,000 cells per well in 24-well ultra-low attachment (ULA) plates (Corning, Incorporated, Corning, NY) in MEBM tumorsphere media (Lonza, Basel, Switzerland) containing 5 µg/mL of insulin (Thermo Fisher Scientific, Waltham, MA), 1 ng/mL of hydrocortisone (Millipore Sigma, Burlington, MA), 2% B27 supplement (Thermo Fisher Scientific, Waltham, MA), 20 ng/mL EGF (Millipore Sigma, Burlington, MA), and 0.1 mM of beta-mercaptoethanol (Millipore Sigma, Burlington, MA). The media was combined with 1% w/v solution of 500 mg methylcellulose (Thermo Fisher Scientific, Waltham, MA) diluted in 50 mL serum-free MEBM, supplemented with the aforementioned factors. Tumorspheres were plated into a 1:1 mixture of tumorsphere media and methylcellulose diluted in MEBM with supplements and grown for 5-7 days at 37°C. Subsequently, cells were infected with MRV (100 PFU/cell) in serum-free MEBM (for 3D conditions). Tumorspheres were counted 7 days post infection. Tumorspheres were either directly counted using a Leica microscope (Wetzlar, Germany) or images were acquired using the All-In-One Fluorescence Microscope BZ-X800 (Keyence, Osaka, Japan), using brightfield at 4X magnification. Images were acquired to capture the entire well, ensuring that all tumorspheres were included. Images were stitched using BZ-X800 Analyzer software. Tumorspheres were counted manually using ImageJ software.

To assess the self-renewal capabilities of the BCSCs, secondary tumorsphere assays were performed. Cells from the primary tumorsphere assays were collected by centrifugation at 900 rpm, washed with 1X Phosphate Buffered Saline (PBS), dissociated by trypsinization, resuspended in stop solution (1X PBS + 25% horse serum), washed with tumorsphere media, and strained using 40-µM Flowmi cell strainers (Corning, Incorporated, Corning, NY). Viable cells were seeded at 1,000 cells per well (n=3 per condition) in 24-well ULA plates in tumorsphere media combined with methylcellulose media (1 mL final volume) as above. The secondary tumorspheres were allowed to grow for 7 days. On day 7, tumorspheres were counted manually and analyzed using the All-In-One Fluorescence Microscope BZ-X800 as described above.

### Serial Passaging

1×10^6^ MCF7 TaxR cells were plated in tumorsphere media and grown for 7 days. Tumorspheres were infected with MRV at 100 PFU/cell in one well of a 6-well ULA plate (Corning, Incorporated, Corning, NY). On the day of infection, fresh TaxR cells were plated in parallel for subsequent infection with passaged virus. After 7 days of infection, 3 freeze/thaw cycles were performed to release the virus. Briefly, TaxR tumorspheres co-cultured with virus (T1L strain) were placed in a container with dry ice and ethanol. The co-culture was incubated for 6-8 minutes until frozen and then placed on the incubator at 37°C until the co-culture was fully thawed. After 3 freeze/thaw cycles, fresh TaxR tumorspheres were pelleted and infected with 1 mL of the freeze/thaw virus/tumorsphere co-culture, fresh tumorsphere media was added, and tumorspheres were incubated at 37°C for 7 days until the next cycle. The serial passaging process was repeated 10 times (once per week for 10 weeks).

### Plaque Assay

Briefly, control or heat-treated virus samples were diluted into PBS supplemented with 2 mM MgCl2 (PBSMg). L929 cell monolayers in 6-well plates (Greiner Bio-One, Kremsmünster, Austria) were infected with 250 μL of diluted virus for 1h at RT. Following the viral attachment incubation, the monolayers were overlaid with 4 mL of serum-free medium 199 (Millipore Sigma, Burlington, MA) supplemented with 1% Bacto agar (BD Biosciences, Franklin Lakes, NJ), 10 μg/mL of TLCK-treated chymotrypsin (Worthington Biochemical, Lakewood, NJ), 2 mM l-glutamine (Invitrogen, Waltham, MA), 100 U/mL of penicillin (Invitrogen, Waltham, MA), 100 μg/mL of streptomycin (Invitrogen, Waltham, MA), and 25 ng/mL of amphotericin B (Millipore Sigma, Burlington, MA). The infected cells were incubated at 37°C, and plaques were counted 5 days post infection.

### Virus Purification

Virus purification of T3D, TIL, and R2 was conducted as previously reported (40). The serial passaged pool (T1L SP) and clones (T1L SP A, T1L SP B, T1L SP C) were similarly purified following plaque assay isolation.

### Western Blot

Cells in 2D conditions were seeded at 2.5 x 10^5^ cells per well in 6-well plates (Corning, Incorporated, Corning, NY) supplemented with 2 mL of media and were incubated for 24h at 37°C. After incubation, MRV was diluted in serum-free media as previously described, but to a final volume of 500 μL. The media was removed, virus was added, and the cells were incubated for 1h at RT rocking. Subsequently, 2 mL of complete media was added, and the cells were incubated at 37°C. The lysates were collected at indicated time points following infection by adding 100-200 μL of RIPA buffer [1M Tris pH: 7.5, 400 mM EDTA, 5 M NaCl, 10% NP-40, 10% SDS, 10% DOC plus a tablet of complete mini protease inhibitor (Millipore Sigma, Burlington, MA) and PhosStop (Millipore Sigma, Burlington, MA)] per 10 mL buffer to each plate and then cells were scraped from the bottom of the plate. RIPA lysates were transferred to a 1.5 mL tube and then centrifuged at maximum speed for 10 minutes. Supernatant was collected in a new tube. Pierce BCA protein assay (Thermo Fisher Scientific, Waltham, MA) was performed for protein quantification according to manufacturer’s instructions. For 3D cultures, tumorspheres were seeded at 2.5-5 x 10^5^ cells per well in 6-well ULA plates (Corning, Incorporated, Corning, NY). Tumorspheres were grown for 5-7 days and infected with MRV (100 PFU/cell) in 500 μL of serum-free media and lysates were collected 24h or 72h post infection. Tumorspheres were collected, washed with 1X PBS, centrifuged, and 100 μL-200 μL of RIPA buffer was added to the cell pellet. Lysates were then processed as described above. Precast gels (Bio-Rad, Hercules, CA) were used and 7-20 μg total protein was loaded per well. Blots were probed with anti-reovirus polyclonal antisera at a 1:1000 dilution for 1h at RT or overnight incubation at 4°C, followed by three washes with 1X PBS with Tween. After this, the secondary goat anti-rabbit HRP antibody (Bio-Rad, Hercules, CA) was used at 1:5000 dilution for 1h at RT followed by three washes with 1X PBS with Tween. Blots were developed using the Super Signal West Pico PLUS Chemiluminescent Substrate (Fisher Scientific, Waltham, MA).

### Flow Cytometry to detect MRV-infected cells

Single cell suspensions were obtained by passing cells diluted in FACS buffer (1XPBS supplemented with 3% FBS) through 40-μM cell strainers, 40-μM Flowmi cell strainers, and a 22G needle with a 3 mL syringe. Cells were counted to obtain 3-5 x 10^5^ for flow analysis. Cells were fixed using the BD CytoFix/CytoPerm fixation/permeabilization solution kit (Fisher Scientific, Waltham, MA) and were incubated with polyclonal reovirus antibody at a concentration of 1:10,000 (2D samples) or 1:2,500 (3D samples) for 1h at RT with rocking. Samples were washed with 1X perm buffer three times. Cells were stained with AlexaFluor (AF) 488 goat anti-rabbit (Invitrogen, Waltham, MA) antibody at 1:2000 for 30 minutes at RT with rocking. Cells were washed with the 1Xperm buffer 2 times, resuspended in 500 μL of 1X perm buffer and transferred to 5 mL round bottom polystyrene tubes (Corning, Incorporated, Corning, NY). The data was acquired using the LSR Fortessa cytometer (BD, Franklin Lakes, NJ) and data analysis was conducted using the FlowJo software.

### RNA sequencing

MCF7 parental and MCF7 TaxR cells were seeded as single cell suspensions (5 x10^5^ cells/well in 6-well ULA plates) for tumorsphere formation (but without methylcellulose). Two wells were combined for each of 3 biological replicates. Tumorspheres were allowed to grow for 7 days. After 7 days, tumorspheres were infected with T1L strain at 100 PFU/cell and RNA was collected at 0, 6 and 24h post-MRV infection using the RNeasy Plus Mini Kit (Qiagen, Germantown, MD). RNA sequencing was conducted by Novogene Corporation, Inc. (Sacramento, CA), using NovaSeq (Illumina, San Diego, CA) generating 6Gb/20M PE150 read-pairs per sample. GEO submission pending.

Differential expression analysis was performed using DESeq2 (v1.44.0) in RStudio. Genes with an absolute log2 fold change ≥ 1 (|log₂FC| ≥ 1) and FDR p-adjusted values (< 0.05) were selected for heatmap visualization to include both upregulated and downregulated genes, followed by retrieval of their normalized counts variance stabilizing transformation (VST) and subsequent z-scored scaling. Venn diagrams were developed by selecting genes from the DESeq2 analysis based on positive and negative log2 fold change values and FDR p-adjusted values (< 0.05). GSEA was conducted using normalized counts with the GSEA software from Broad Institute v4.3.3.

### Viral RNA sequencing

Viral RNA was extracted by taking 10 μL of purified virus and mixing with 90 μL of TriPure Isolation Reagent.(Millipore Sigma, Burlington, MA). A 200 μLvolume of chloroform (Millipore Sigma, Burlington, MA) was added to the sample and vortexed until phase separation occurred, forming a clear upper aqueous layer. After centrifugation (13,000 rpm for 10 min at 4°C), the upper layer (containing RNA) was transferred to a new tube. RNA was precipitated with 600 μL isopropanol, was left incubating for 10 minutes at RT, pelleted by centrifugation (13,000 rpm for 10 minutes at RT), washed with 75% ethanol and pelleted by centrifugation (13,000 rpm for 5 minutes at RT) Subsequently, the supernatant was removed and the pellet was left to air dry in a heat block at 42°C and resuspended in RNase-free water. RNA sequencing was conducted by the Center for Genomics and Bioinformatics at Indiana University, Bloomington.

Libraries were prepared by Takara (Clontech) SMARTer Stranded RNA-Seq Kit protocol (catalog no. 634837). Briefly, a modified N6 primer primes the first-strand synthesis reaction. When the reverse transcriptase reaches the 5′ end of the RNA, its terminal transferase activity adds a few non-templated nucleotides. A specialized SMARTer (Switching Mechanism at the 5′ end of RNA Template) oligonucleotide base-pairs with these added nucleotides, acting as an extended template, allowing the reverse transcriptase to continue synthesizing full-length, single-stranded cDNA. The purified first-strand cDNA is PCR amplified into RNA-seq libraries incorporating Illumina indexes. The amplified library is purified using AMPure XP beads, then analyzed by Agilent 4200 TapeStation. Libraries were sequenced on an Illumina NextSeq 1000/2000 P2 Reagents (100 Cycles) v3 flow cell (catalog no. 20046811) configured to generate paired end reads. The demultiplexing of the reads was performed using bcl2fastq, version 2.20.0. GEO submission pending.

### Real Time Glo Extracellular ATP (eATP) assay

Real Time Glo Extracellular ATP (eATP) assay was purchased from Promega (Madison, WI). Cells were seeded at 5,000 cells per well in white opaque 96-well plates (Corning, Incorporated, Corning, NY). As previously described in the cell viability assay methods section, after 24h of plating, media was removed and cells were infected with MRV strains at 100 PFU/cell diluted in serum-free media (25 μL) for 1h at RT with rocking and subsequently, 100 μL complete media was added. Cells were incubated at 37°C for 4 days. After 4 days of incubation, eATP reagent was prepared at a concentration of 4X and 50 μL of the reagent was added to each well. Luminescence readings were recorded using the BioTek Synergy 2 Microplate Reader along with the Gen5 software (BioTek Instruments, Inc, Winooski, VT)

For inhibitor studies, cells were seeded and infected as described above, but using a final volume of 12.5 µL. Twenty-four hours post infection, Q-VD-Oph (20 µM) (SelleckChem, Houston, TX) was added at 12.5 µL per well, followed by incubation at 37°C for 48h, cells were then retreated with the same inhibitor concentration, and eATP was measured on the following day.

### Immunofluorescence

Cells were plated at a density of 5,000 cells per well in 4-well chamber slides (Thermo Fisher Scientific, Waltham, MA) and infected (100 PFU/cell) after 24h. Cells were fixed with 4% PFA, then permeabilized for 30 minutes on ice (1X PBS + 0.5% Triton X-100), blocked (200 mL 1X PBS + 0.1% BSA + 0.2% Triton X-100 + 0.05% Tween) and μNS primary antibody was added at 1:1000 for 1h. Fixed cells were washed 3X with 1X PBS and AlexaFluor (AF) 488 goat anti-rabbit (Invitrogen, Waltham, MA) was added at 1:500, for 1h. Fixed cells were washed 3X with 1X PBS, DAPI (5 mg/mL stock) was added at 1:5000 in 1X PBS, and washed twice with 1X PBS. The cells were mounted using Prolong Diamond (Thermo Fisher Scientific, Waltham, MA) and imaged after 24h using the All-In-One Fluorescence Microscope BZ-X800.

### Live/Dead Cell Imaging Kit (488/570)

Live/Dead Cell Imaging Kit (488/570) was purchased from Thermo Fisher Scientific. MCF7 and TaxR tumorspheres were grown for 7 days in 96-well ULA plates (Corning, Incorporated, Corning, NY) as previously described in the tumorsphere assay methods section. After 7 days, cells were infected with the MRV strains and incubated for 7 days. After 7 days, the live/dead reagents were mixed at a 1:1 ratio and added to the media for 15 minutes according to the manufacturer’s instructions. Every well was captured and stitched using the All-in-One Fluorescence Microscope BZ-X800. Quantification was conducted based on the number of live (green) and dead/dying (red) tumorspheres.

### qRT-PCR

Tumorspheres were plated as previously described. Tumorspheres were collected, washed with 1X PBS, and RNA was collected using TriPure following the manufacturer’s protocol. RNA quantity and purity (260/280) were assessed using the Take 3 Plate on the BioTek Synergy 2 Microplate Reader, and 500 ng of total RNA per sample was used for cDNA synthesis. cDNA was prepared using the appropriate amount of RNA (500 ng-1000 ng), qSCRIPT (Quantabio, Beverly, MA) and DEPC water (Santa Cruz Biotechnology, Dallas, TX) to a final volume of 10 µL according to the manufacturer’s instructions. Standard curve samples of cDNA were prepared using serial dilutions of 1:1, 1:5, 1:10, 1:20, 1:40 and 1:80 in DEPC water. The qPCR was performed using SYBR Green master mix (Roche, Basel, Switzerland) in a 10 µL reaction volume containing cDNA, gene-specific primers, and RNase-free water (Qiagen, Germantown, MD) in LightCycler 480 qPCR plates (Roche, Basel, Switzerland). Gene expression was calculated using the standard curve, and the 18S rRNA gene as an internal control. Z-score values were calculated across genes, n=3.

### Statistical Analysis

Cell viability assays, primary and secondary tumorsphere assays, flow cytometry, immunofluorescence, eATP assay and inhibitor studies were analyzed using Two-Way ANOVA. For comparisons of two groups, unpaired T Test was conducted. Live/Dead assay results for each cell line (Figure 4) were analyzed using Ordinary One-Way ANOVA. Statistical significance is indicated as follows: *p ≤ 0.05, **p ≤ 0.01, ***p ≤ 0.001, and ****p ≤ 0.0001. The data was analyzed using GraphPad Prism. Experiments were repeated at least three times.

## Results

### Effects of MRV strains on 2D and 3D cultures

To explore how MRV strains affect the cell viability in a panel of therapy sensitive and therapy resistant ER+ breast cancer cells (**Table 1)** we performed alamarBlue cell viability assays. Under standard 2D growth conditions, cells were exposed to either mock (serum-free media) or 3 different strains of MRV: T3D, T1L and R2. Four days post infection, alamarBlue was added to each well to assess viability. All MRV strains decreased cell viability in all cell lines compared to mock-treated cells. No cell line-specific differences were observed (**Figure 1A**), with the exception of the non-transformed MCF10A cells. We found that T3D decreased cell viability in MCF10A cells, but T1L and R2 did not, indicating that T3D may have a killing effect in non-transformed mammary epithelial cells. For further studies, we focused on examining differences between parental MCF7 and paclitaxel-resistant (TaxR) MCF7 cells (herein referred to as TaxR) as we have found that TaxR cells are enriched for BCSC populations **(Supplementary Figure S1 A-C)**. Intracellular staining with an anti-reovirus polyclonal antisera followed by flow cytometry was used to quantify the percentage of infected cells in 2D conditions. Overall, parental MCF7 cells had a higher number of cells infected compared to TaxR cells, particularly following T3D infection (**Figure 1B**). To determine if viral protein synthesis is different between MCF7 parental and TaxR cells, western blotting was performed on protein lysates collected from cells 24h post infection with T3D, T1L or R2. Similar levels of the outer capsid protein, μ1, were observed in all infected lysates in MCF7 and TaxR cells (**Figure 1C**). Next, we assessed the effect of MRV strains on extracellular ATP (eATP), a DAMP (damage-associated molecular pattern) released upon oncolytic virus infection and a marker of immunogenic cell death (41,42). We tested the levels of eATP in the media from MCF7 and TaxR cells treated with mock, T3D, T1L and R2. Interestingly, we found that T1L and R2 induce higher levels of eATP compared to T3D in both MCF7 and TaxR cells (**Figure 1D**). In summary, under 2D conditions, we observed that parent MRV strains are efficient at decreasing cell viability, infecting cells, and producing viral proteins, but T1L and R2 may be more efficient inducers of immunogenic cell death.

**Figure 1.**
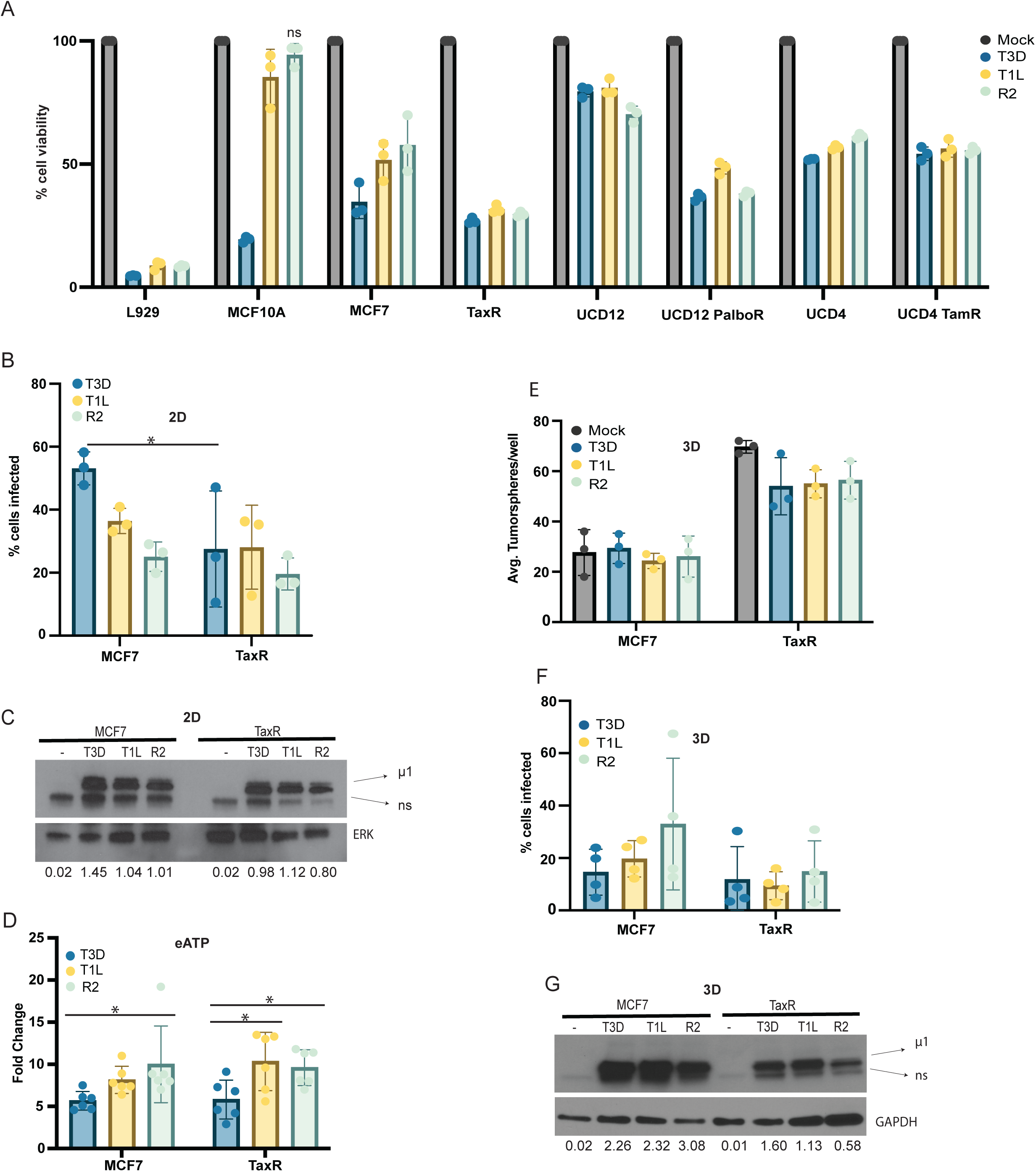
Parent MRV strains show high oncolytic effects in 2D conditions but limited effects in BCSCs. **A)** alamarBlue assay 4 days post infection. Cell viability was calculated by dividing infected cell alamarBlue readings by mock infected cells x 100 (percent viability), n=3. All comparisons of parent MRV to mock were statistically significant <0.0001, with the exception of R2 in MCF10A cells. **B)** Flow cytometry was used to quantify the percentage of cells infected using a polyclonal reovirus antibody 24h post infection in 2D conditions, n=3. **C)** Western blotting of reovirus proteins using the reovirus antibody, 24h post infection in 2D conditions. Values correspond to normalization to ERK loading control. **D)** Extracellular ATP assay (eATP) assay, 4 days post infection. Luminescence values were normalized to mock samples, n=6. **E)** Tumorsphere numbers 7 days following infection with parent viruses, n=3. **F)** Quantification of tumorspheres 24h post infection using flow cytometry, n=4. **G)** Western blotting using reovirus antibody 24h post infection in 3D conditions. Values correspond to normalization to GAPDH loading control. All statistical analysis was conducted using Two-Way ANOVA. Statistical significance is indicated as follows: *p ≤ 0.05, **p ≤ 0.01, ***p ≤ 0.001, and ****p ≤ 0.0001.

**Table 1.** Summary of cell lines tested for sensitivity to MRV.

| Cell line | Description |
| --- | --- |
| L929 | murine fibroblasts |
| MCF10A | human mammary epithelial cells, non-transformed |
| MCF7 | ER+, parental cell line |
| TaxR | ER+, paclitaxel resistant |
| UCD12 | ER+, patient-derived xenograft (PDX) cell line |
| UCD12 PalboR | ER+, PDX cell line, palbociclib resistant |
| UCD4 | ER+, PDX cell line |
| UCD4 TamR | ER+, PDX cell line, tamoxifen resistant |

Next, we aimed to better understand the effects of MRV on BCSCs. MCF7 and TaxR cells were cultured as 3D tumorspheres in serum-free conditions in ultra-low attachment plates (ULA) to enrich for BCSCs, as we have previously published (38). Tumorspheres were grown for 5-7 days prior to MRV infection (mock, T3D, T1L or R2) and then incubated for 7 days. Tumorspheres per well were then counted to assess the effect of MRV strains on tumorsphere killing. We found that all MRV strains have a minimal effect on killing tumorspheres compared to mock infection for both MCF7 parental and TaxR tumorspheres (**Figure 1E**). As with 2D infected samples, flow cytometry was performed to determine the percentage of cells infected by MRV in 3D tumorspheres **(Figure 1F)**. Parental MCF7 cells tended to have a higher number of cells infected compared to TaxR cells, though it did not reach statistical significance. Additionally, the percentage of cells infected in 3D tumorsphere conditions was generally lower compared to 2D conditions for both MCF7 parental and TaxR cells (**Figure 1B, F**). Viral protein synthesis was assessed in tumorspheres that were infected with mock, T3D, T1L or R2 for 24h prior to collecting protein lysates. We observed lower levels of the μ1 protein in the TaxR tumorspheres compared to the MCF7 parentals (**Figure 1G**). In summary, when cells are cultured in 3D conditions, the parent MRV strains have modest effects in reducing tumorspheres, as well as lower infection efficiency and viral protein synthesis.

### TaxR tumorspheres exhibit a higher interferon response at baseline and post infection

The molecular mechanisms of MRV infection and cell death in ER+ BCSC populations are unknown. Although T3D is the most extensively studied MRV strain, we found it significantly reduced cell viability in MCF10A normal mammary epithelial cell line. Furthermore, the T1L strain, which remains relatively understudied in the context of MRV OVs, was associated with higher levels of eATP, indicating more cell damage and a pro-inflammatory signature. Several studies have shown that innate immune responses to T1L are context-dependent (43,44). Based on these distinct observations, we opted to focus our studies on the effects of T1L on BCSCs. To elucidate the effects of MRV in BCSCs, we leveraged RNA sequencing using MCF7 and TaxR cells grown as tumorspheres. Tumorspheres were infected with T1L, and RNA was collected at baseline (mock) and then 6– and 24-h post infection. Principal Component Analysis (PCA) showed that MCF7 parental and TaxR samples cluster separately at all time point conditions, which indicates significant transcriptomic differences between these cell lines, including at baseline (mock) (**Figure 2A**). Using differential expression gene analysis (DESeq2), we extracted relevant genes based on FDR p-adjusted values and absolute log2 fold change ≥ 1 (|log₂FC| ≥ 1) heatmap visualization to include both upregulated and downregulated genes, followed by retrieval of their normalized counts variance stabilizing transformation (VST) of MCF7 and TaxR 24h/mock. Genes were subsequently z-scored for downstream visualization. (**Figure 2B**). Based on the z-score analysis, TaxR cells have upregulation of genes related to antiviral defenses such as IFIT1, IFIT2, IFIT3, OAS1, APOL1, APOL2, ZC3HAV1, RIG-I, ISG15, DDX60 and others at baseline compared to MCF7 and 24h post infection (**Figure 2B**). From the differential gene expression analysis, genes meeting the criteria of significant FDR-adjusted p-values (p<0.05) and both positive and negative log2 fold change values were selected. Based on Venn diagrams, we found 87 shared upregulated genes and 55 shared downregulated genes at 24h post infection compared to mock conditions (**Figure 2C**), reflecting 9.6% and 10.9 % of genes, respectively. Pathway analyses using EnrichR (45) (Reactome and MSigDB) was performed on both the overlapping and uniquely up– and downregulated genes. We observed distinct Reactome pathways that were suppressed and activated (**Figure 2D**). Shared activated pathways included viral response pathways, interferon signaling, cytokine signaling, and NF-κB activation though FADD. Downregulated or suppressed shared pathways included rRNA processing and modification, SUMOylation of transcription co-factors, B-WICH that positively regulated rRNA expression, as well as other pathways (**Figure 2D, Supplemental tables**). The unique MCF7 upregulated and downregulated genes were also entered into EnrichR and found to be associated with antiviral response and RNA pol I transcription Reactome pathways, respectively (**Supplementary Figure S2**). The TaxR unique upregulated genes were associated with cell cycle and Rho GTPases Reactome pathways, and downregulated genes were associated with translation (**Supplementary Figure S2**). Upon examining pathways cataloged in MSigDB (46), we determined that shared upregulated genes were associated with cellular stress, immune signaling, and inflammation, while shared downregulated genes were linked to proliferation, growth, and metabolism (**Supplementary Figure S3**). In MCF7 cells, unique upregulated pathways included immune signaling, inflammation, stress responses, and cell cycle regulation, with downregulated pathways related to proliferation, metabolism, and cell cycle control (**Supplementary Figure S3**). In TaxR cells, uniquely upregulated pathways included cell cycle regulation, proliferation, immune signaling, and developmental programs, whereas downregulated pathways reflected growth, metabolism, proliferation, and stress responses (**Supplementary Figure S3**). The full gene lists and EnrichR results for Reactome and MSigDB Hallmark pathways are included in **Supplementary Tables 2 and 3 and Supplementary Figures S2 and S3.** Gene set enrichment analysis (GSEA) was performed for MCF7 parental and TaxR comparisons (6h vs. mock and 24h vs mock). GSEA hallmark pathways visualized using a bubble plot show that in both MCF7 and TaxR tumorspheres interferon alpha and interferon gamma pathways were significantly upregulated at 24h post infection (**Figure 2E**), In TaxR tumorspheres, there was a decrease in interferon pathways at 6h; however, at 24h there was a significant activation (**Figure 2E, F**). Overall, this data indicates that in MCF7 tumorspheres, T1L induces an innate immune response (i.e. interferon alpha and gamma) earlier than in the TaxR tumorspheres, but by 24h, the TaxR response is comparable to the MCF7 cells. Of note, GSEA of mock infected MCF7 and TaxR tumorspheres indicates that TaxR tumorspheres have a higher interferon/inflammatory response pathway activation at baseline (**Figure 2E**), consistent with lower infection rates in the TaxR cells, as a strong antiviral response suppresses viral replication. Further, the activation of these pathways suggests immune response activation in response to MRV.

**Figure 2.**
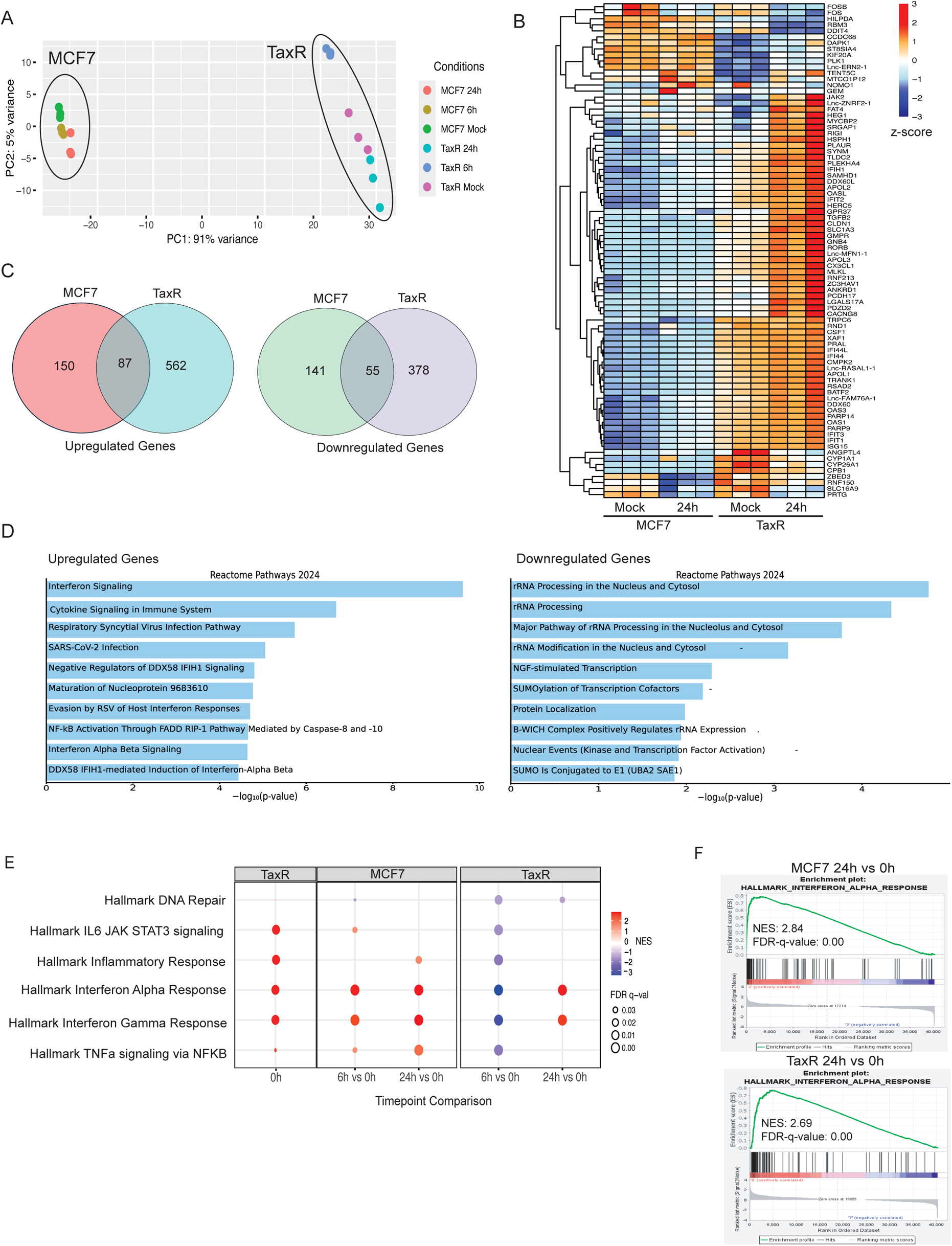
TaxR tumorspheres exhibit a higher interferon response 24 hours post infection. **A)** Principal Component Analysis (PCA) of MCF7 and TaxR cells grown as tumorspheres, infected with T1L 0h (mock), 6h, and 24h post infection, n=3. **B**) Heatmap of z-scored normalized counts VST generated based on selecting genes with absolute log2 fold change ≥ 1 (|log₂FC| ≥ 1) to include both upregulated and downregulated genes and significant FDR p-adjusted values (<0.05). **C)** Venn diagrams were generated by selecting genes from the DESeq2 analysis based on all positive and negative log2 fold change values and FDR p-adjusted values (<0.05) in both cell lines 24h post infection. **D)** Overlapping genes from the Venn diagrams were entered into EnrichR (Reactome) for downstream pathway analysis to obtain upregulated and downregulated pathways at 24h post infection**. E)** Gene Set Enrichment Analysis (GSEA) Hallmark Pathways significantly up or downregulated based on the Normalized Enrichment Score (NES) and FDR q-value. **F)** Interferon alpha pathway shows positive NES scores in both BCSCs at 24h vs mock.

### Serial passaging T1L strain results in a superior oncolytic virus

As MRV strains T3D, T1L and R2 did not have a robust killing effect on MCF7 parental and TaxR tumorspheres, we aimed to develop a selective MRV strain with enhanced capacity to kill BCSCs. We focused on further developing T1L through serial passaging (**Figure 3A**) because this strain is known to be less sensitive to interferon signaling and BCSCs have been shown by multiple groups to have high interferon signaling (47–52). Here, TaxR cells were used for serial passaging due to their enrichment for BCSC-like characteristics such as high ALDH1 activity, increased CD44^hi^ /CD24^lo^ populations, and higher tumorsphere formation capability (**Supplementary Figure S1 A-C**). For serial passaging, T1L was allowed to infect and replicate for 7 days in TaxR tumorspheres, virus-containing supernatant was collected by 3 freeze/thaw cycles, and then the passaged virus was infected into intact fresh TaxR tumorspheres. After 10 passages, there was a presumed mixture of multiple viral strains, denoted as T1L SP (serial passaged) pool (**Figure 3A**). We hypothesized that, being adapted to TaxR tumorspheres, T1L SP would exhibit enhanced oncolytic potential. Cell viability assays showed that T1L SP significantly decreased cell viability more effectively than T1L in MCF7 parental and TaxR cells cultured in 2D conditions (**Figure 3B**). In 3D tumorsphere assays, T1L SP significantly reduced tumorsphere formation in both MCF7 parental and TaxR tumorspheres but the effect was greater in TaxR tumorspheres (**Figure 3C**). Secondary tumorsphere assays can assess self-renewal capacity of BCSCs and can be used to determine how many BCSCs survive MRV treatment. After the primary tumorsphere assay, tumorspheres were dissociated, replated, grown for 5-7 days, and counted. In the secondary tumorsphere assays, the effect of T1L SP was more robust, suggesting fewer BCSCs survived compared to treatment with T1L (**Figure 3D**). UCD4 parental and UCD4 TamR (tamoxifen resistant) cell lines derived from the UCD4 PDX model were included in these studies to determine if this effect was specific to the MCF7 TaxR model. Interestingly, the results showed that T1L SP does reduce tumorsphere formation more effectively than T1L in both UCD4 parental and TamR tumorspheres (**Supplementary Figure S4A**). A Live/Dead Imaging assay was used to determine if T1L SP causes more cell death than T1L (**Figure 3E**). The results demonstrated that T1L SP induces significantly greater cell death than T1L in both cell lines (**Figure 3E**). Representative images of T1L and T1L SP can be found in **Supplementary Figure S4B**. Overall, these findings suggest that T1L SP is more potent than T1L, with its effects on cell viability, tumorsphere formation, and cell death most notable in the TaxR cells.

**Figure 3.**
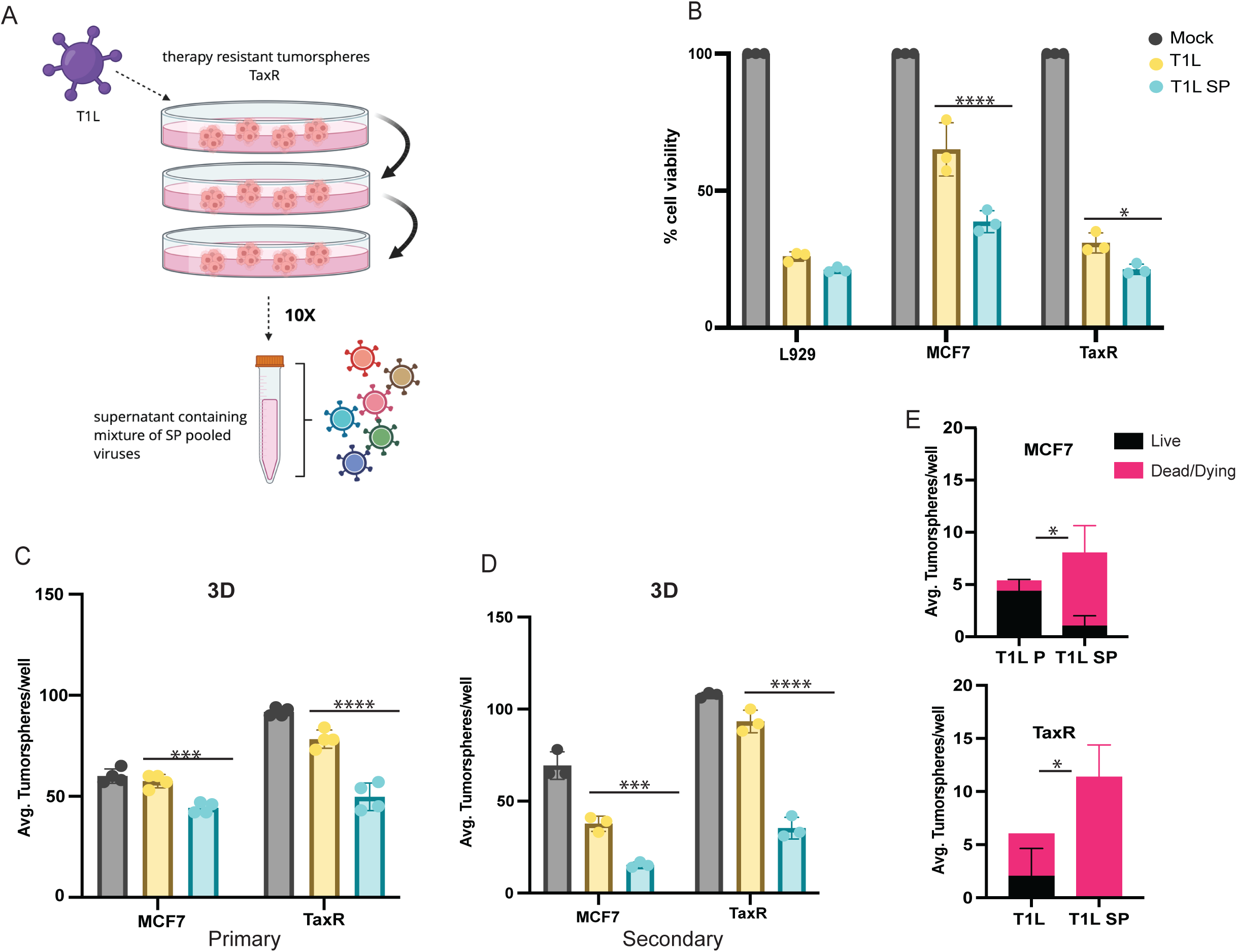
Serial passaging T1L strain results in a superior oncolytic virus. **A)** Schematic of the development of serial passaged pool viruses in therapy-resistant TaxR tumorspheres using T1L. **B)** alamarBlue assay was used to determine the effect of T1L and T1L serial passaged (SP) pool in 2D conditions at 4 days-post infection, n=3, Statistical Analysis: Two-Way ANOVA. **C).** MCF7 and TaxR were grown as tumorspheres to determine the effect of T1L and T1L SP, Statistical Analysis: Two-Way ANOVA, n=4. **D)** The tumorspheres shown in (C) were dissociated and replated to assess the survival and self-renewal capacity of the treated cells (secondary tumorsphere assay), n=3, Statistical Analysis: Two-Way ANOVA. **E)** Quantification of tumorspheres infected with T1L or T1L SP using the Live/Dead Imaging Kit, n=3, Statistical Analysis: Unpaired Student’s T Test. Statistical Analysis: Unpaired T Test. Statistical significance is indicated as follows: *p ≤ 0.05, **p ≤ 0.01, ***p ≤ 0.001, and ****p ≤ 0.0001.

### Serial Passaged MRV clones have differential effects in BCSCs

As T1L SP likely contains a pool of different T1L variants with different oncolytic properties, we aimed to isolate specific viral clones from the pool of serial passaged viruses. T1L SP viral strains were isolated using a plaque assay and three different clones (T1L SP A, T1L SP B, T1L SP C) were randomly selected and purified (**Figure 4A**). The clones were tested for their cell killing properties in comparison to T1L. In 2D cell viability assays, the clones did not decrease cell viability as robustly as T1L (**Figure 4B**). We assessed cell death in tumorsphere conditions using the Live/Dead Imaging Kit, and we observed an increase in dead/dying tumorspheres with the MRV clones, compared to T1L in TaxR tumorsphere cultures (**Figure 4C, D**). The dead/dying spheres were much higher in T1L SP B and T1L SP C in the TaxR tumorspheres compared to the MCF7 tumorspheres (**Figure 4C, D**). This data indicates that T1L SP B and T1L SP C selectively kill therapy-resistant TaxR tumorspheres. To determine if the increase in tumorsphere death is the result of an increase in the percentage of cells infected, we performed flow cytometry. MCF7 parental and TaxR tumorspheres were infected with T1L, T1L SP A, T1L SP B, and T1L SP C for 24h and then single cells were stained with an MRV-specific antibody prior to flow cytometry. Interestingly, the percentage of infected cells was higher in T1L infected spheres compared to the clones in both MCF7 and TaxR tumorspheres (**Figure 4E**). Western blots showed that T1L produced more viral protein compared to T1L SP A, T1L SP B, and T1L SP C in infected MCF7 tumorsphere lysates at 24h post infection (**Figure 4F**). T1L SP B and T1L SP C produced similar levels of μ1 protein and T1L SP A produced the least amount of μ1 protein in MCF7 tumorspheres (**Figure 4F**). Similar trends were observed in TaxR tumorspheres, whereby T1L produced abundant levels of μ1 protein compared to the other clones at 24h post infection (**Figure 4F**). At 72h post infection, we observed more expression of μ1 in T1L SP B and T1L SP C, especially in the TaxR tumorspheres (**Figure 4F**), possibly indicating a difference in viral protein synthesis kinetics between T1L, T1L SP B, and T1L SP C. To assess viral replication over time, an increase in virus titer relative to the input from MCF7 parental and TaxR tumorspheres infected with T1L, T1L SP B and T1L SP C over 6 days was compared. For these experiments, virus titer in the media (supernatant) of cells at each time point was quantified by plaque assay. Whereas no difference in virus replication efficiency between strains were observed in MCF7 tumorspheres, an increase in infectious virus production was seen in the T1L SP B TaxR infected tumorspheres compared to T1L, indicating better replication efficiency and better adaptation (**Figure 4G**). Together, these data suggest that the SP clones have adapted for enhanced viral replication and subsequent killing in the TaxR tumorspheres.

With the aim of identifying acquired mutations during serial passaging, the viral RNA from the SP clones was sequenced. Several mutations were identified, including amino acid changes in proteins that play a role in attachment (e.g. σ1) and viral replication (e.g. µNS), see **Table 2**. We focused on studying T1L SP B and T1L SP C, due to their potent cell killing effects (**Figure 4D**). T1L SP B and T1L SP C contain one point mutation in the μNS protein, which is a nonstructural protein, encoded by the M3 segment that forms viral factories and interacts with other nonstructural and structural proteins to coordinate genome replication. We hypothesized that the mutations in μNS may facilitate the formation of viral replication factories. We aimed to assess viral factories via μNS immunofluorescence. We visualized μNS positive cells in MCF7 and TaxR cell lines at 24h, 48h and 72h post infection. The μNS positive cells were quantified and divided by the total number of DAPI cells to obtain the percentage of μNS positive cells. Over time, the percentage of μNS-positive cells increased in the T1L SP B (24h vs 72h; p < 0.0001) and T1L SP C (24h vs 72h; p < 0.01), whereas the μNS-positive cells remain constant or decrease over time in cells infected with T1L (**Supplementary Figure S5 A, B**). In MCF7 cells, we quantified more μNS-positive cells in T1L SP B (24h vs 48h; p < 0.05) and T1L SP C (24h vs 48h; p < 0.01) infected cells at 48h post-infection. (**Supplementary Figure S5 A, B**). Together, the μNS immunofluorescence, western blots (**Figure 4F**), and plaque assay (**Figure 4G**) data suggest that T1L SP B and T1L SP C display enhanced viral replication in TaxR tumorspheres.

**Table 2:** Summary of mutations found in serial passaged MRV clones.

| Virus | Mutations | Base-pair change |
| --- | --- | --- |
| T1L SP A | N94D in $\sigma 1$<br>silent mutation in $\sigma 1s$<br>P707S in $\mu NS$<br>silent mutation in $\mu 2$<br>silent mutation in $\lambda 1$<br>silent mutation in $\lambda 1$<br>silent mutation in $\lambda 2$<br>P503S in $\lambda 2$ | S1 A<G position 293<br>S1 A<G position 293<br>M3 C<T position 2119<br>M1 T<C position 1480<br>L3 C<T position 2059<br>L3 G<C position 2062<br>L2 A<T position 90<br>L2 C<T position 1507 |
| T1L SP B | P707S in $\mu NS$<br>silent mutations in $\lambda 1$<br>silent mutations in $\lambda 1$<br>silent mutation in $\lambda 2$ | M3 C<T position 2119<br>L3 C<T position 2059<br>L3 G<C position 2062<br>L2 A<T position 90 |
| T1L SP C | L488S in $\mu NS$<br>silent mutations in $\lambda 1$<br>silent mutations in $\lambda 1$<br>silent mutation in $\lambda 2$ | M3 T<C position 1483<br>L3 C<T position 2059<br>L3 G<C position 2062<br>L2 A<T position 90 |

### Parent MRV and clones both trigger apoptosis, but clones elicit greater immunogenicity

To further understand how SP clones induce cell death, we performed a series of experiments testing the effect of distinct cell death inhibitors to elucidate the dependence of MRV on cell death pathways. For these studies, we focused on T1L SP B and T1L SP C, as T1L SP A showed limited viral production and poor infectivity. Apoptosis is the most frequently observed type of cell death induced by MRV (53–55). We treated MCF7 and TaxR with the pan-caspase inhibitor Q-VD-Oph with and without MRV (T1L, T1L SP B or T1L SP C). We found that the pan-caspase inhibitor resulted in a statistically significant increase in cell viability in the presence of MRV in all cases (**Figure 5A-C**). Overall, this data suggests that T1L, T1L SP B and T1L SP C induce apoptotic cell death. However, because we did not observe a complete inhibition of MRV-induced cell death, and MRV has been shown to induce necrotic cell death in L929 cells that is dependent on the activity of RIPK1 (56), we also assessed the effects of RIPK1 kinase inhibitor, necrostatin-1. In MCF7 parental cells, necrostatin-1 blocked the effect of all MRV clones, but in the TaxR cells, it only blocked the effect of T1L (**Figure 5D-F**). This data shows that apoptosis and necrosis collectively contribute to T1L-induced cell death in both cell lines. Additionally, this data suggests that the cell lines respond differently to the clones, particularly in the case of TaxR cells, where clone T1L SP B and T1L SP C seem to elicit more apoptosis than necroptosis. Dual treatment of necrostatin-1 and Q-VD-Oph with T1L, T1L SP C or T1L SP B also resulted in an increase in cell survival (**Supplementary Figure S6 A-C**) but not greater than with Q-VD-Oph alone.

**Figure 4.**
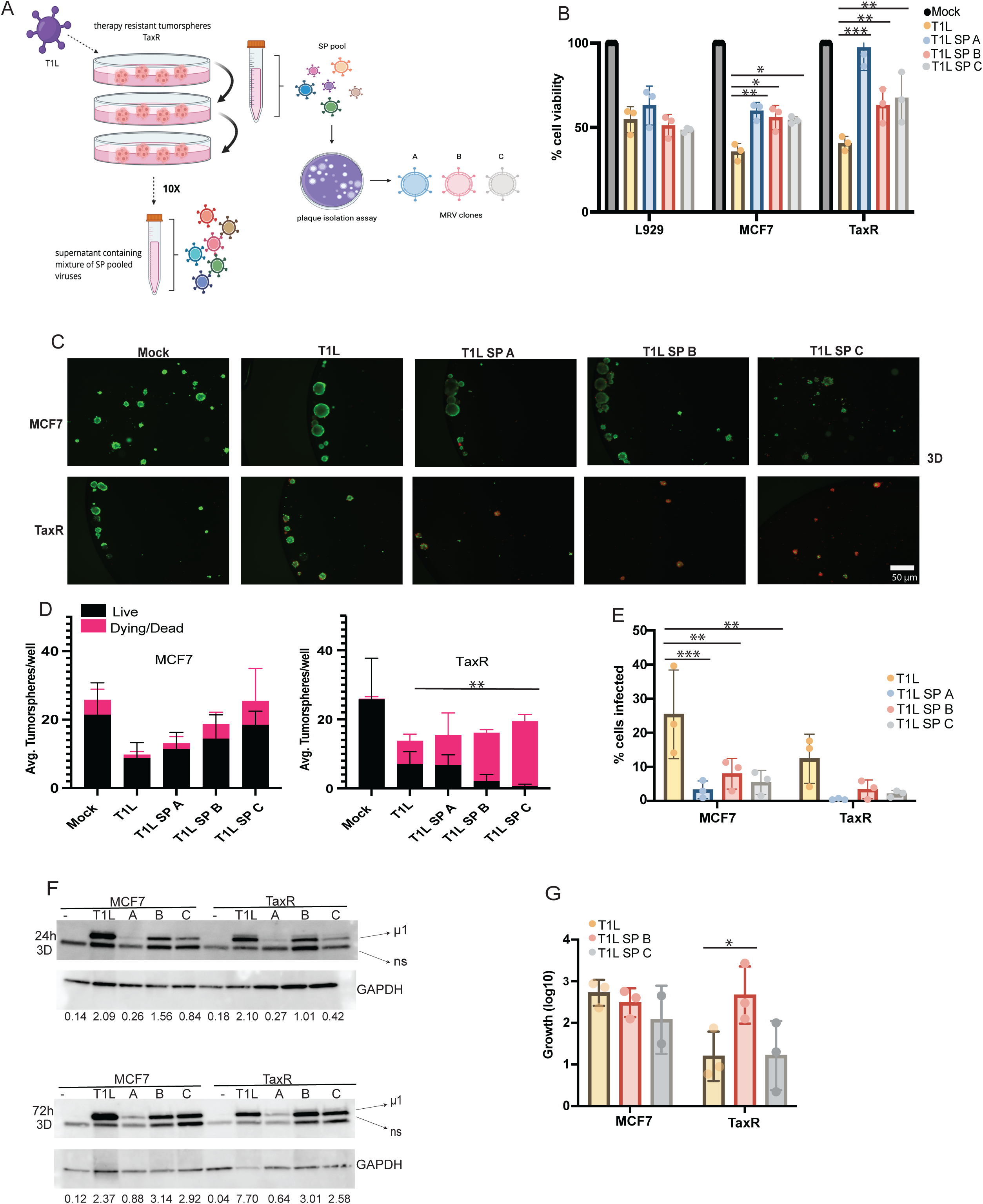
Isolated MRV clones have differential effects in BCSCs. **A)** Schematic depicting the development of viral clones from the serial passaged pool. **B)** alamarBlue assay in 2D conditions was used to assess the effect of viral clones in comparison with T1L, n=3, Statistical Analysis: Two-Way ANOVA. **C)** Live/Dead Imaging Kit was used to assess the viability of tumorspheres infected with T1L and the viral clones. **D)** Quantification of images from (C), n=3, Statistical Analysis: Ordinary-One-Way ANOVA. **E)** Flow cytometry was used to determine the percentage of cells infected with T1L and the SP clones, 24h post infection, n=3. Statistical Analysis: Two-Way ANOVA. **F)** MCF7 and TaxR cells were cultured as tumorspheres and infected for 24h and 72h with T1L, T1L SP A, T1L SP B, and T1L SP C and Western blotting was used to detect viral proteins. Values represent normalization to GAPDH. **G)** Plaque assay of tumorspheres (supernatant and tumorspheres) at day 6 post infection (T1L, T1L SP B, T1L SP C) normalized to 6h post infection, n=3, Statistical Analysis: Two-Way ANOVA. Statistical significance is indicated as follows: *p ≤ 0.05, **p ≤ 0.01, ***p ≤ 0.001, and ****p ≤ 0.0001.

**Figure 5.**
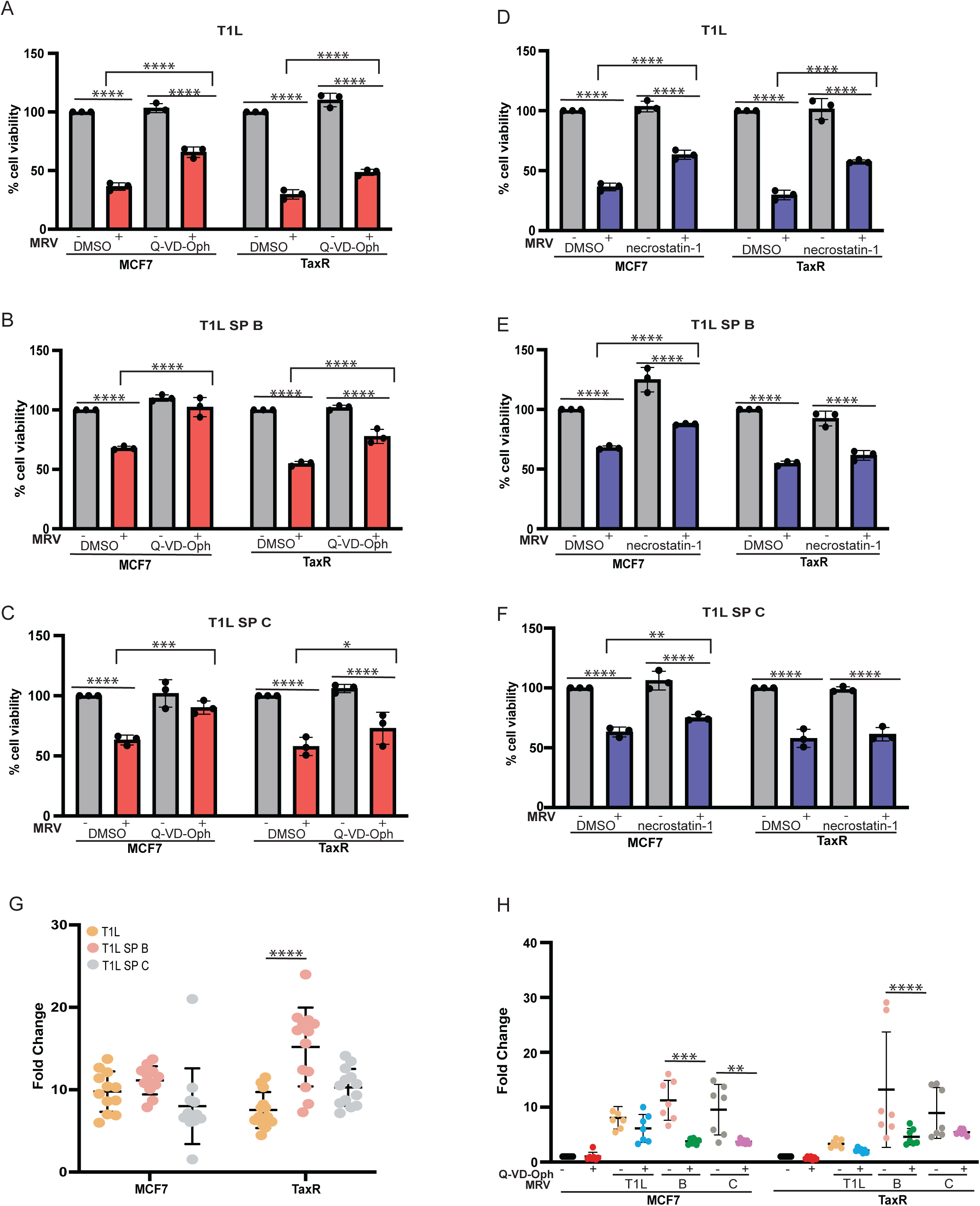
T1L parent and clones both trigger apoptosis, but clones elicit greater immunogenicity. **(A-C)** MCF7 and TaxR cells were plated in 2D conditions infected with: **A)** T1L P, **B)** T1L SP B, or **C)** T1L SP C, and treated with DMSO or apoptosis inhibitor Q-VD-Oph at 20 μM, n=3, **(D-F)** MCF7 and TaxR cells were infected with: **D)** T1L **E)** T1L SP B or **F)** T1L SP C and treated with DMSO or necroptosis inhibitor necrostatin-1 at 50 μM, n=3. **G)** eATP levels were measured 4 days post infection in cells infected with T1L, T1L SP B, or T1L SP C. Data was normalized to the mock control, n=13. **H)** eATP levels were measured 4 days post infection in cells infected with T1L, T1L SP B or T1L SP C with or without Q-VD-Oph and DMSO control was used for data normalization, n=7. All statistical analysis was conducted using Two-Way ANOVA. Statistical significance is indicated as follows: *p ≤ 0.05, **p ≤ 0.01, ***p ≤ 0.001, and ****p ≤ 0.0001.

Given that T1L induced more eATP in TaxR cells (**Figure 1D**) and the RNA sequencing data revealed that interferon alpha (IFN-α) and interferon gamma (IFN-γ) were activated in both cell lines, we hypothesized that the T1L SP clones may also stimulate more immunogenic cell death. MCF7 and TaxR cells were infected with T1L, T1L SP B and T1L SP C for 4 days prior to eATP measurement. We observed that in the TaxR cells, there was a significant increase in the levels of eATP in T1L SP B compared to T1L (**Figure 5G**). Interestingly, treatment with Q-VD-Oph significantly reduced eATP release induced by T1L SP B in the TaxR cells, and T1L SP B and T1L SP C induced eATP release in the MCF7 cells, indicating that the immunogenic cell death is dependent on apoptosis (**Figure 5H**).

### The viral clones elicit weaker antiviral responses compared to the parental T1L strain

To assess whether T1L SP B and T1L SP C differed from the parental T1L strain in their antiviral response, we performed qRT-PCR for the presence of the L1 segment of T1L, and IFIT1, which we found differentially regulated in RNA-seq studies (**Figure 2B**). The L1 gene segment, which encodes for viral core protein λ3, as a marker for viral replication. RNA was collected from MCF7 TaxR tumorspheres at 24, 72, and 96h post infection. As shown **in Figure 6A**, T1L infection induces sustained expression of the L1 segment following infection, while expression of L1 in T1L SP B and T1L SP C infected cells increases over time, only reaching the same level of T1L infected cells at 96 hours. As we observed in Figure 2, IFIT1 expression is induced within 24h of infection with T1L. Interestingly, we found that induction of IFIT1 expression is not as robustly induced with T1L SP B and T1L SPC (**Figure 6A**). We expanded our qRT-PCR to a handful of selected key antiviral genes from the RNA-seq comparison. At 24h post infection, T1L induced a noticeably stronger antiviral response, reflected by higher expression of IFIT1, IFIT2, IFIT3, RIG-I, and IFIH1. T1L infection resulted in a time-dependent reduction in antiviral gene expression, whereas T1L SP B and T1L SP C clones displayed a modest, time-dependent increase in expression of these same genes (**Figure 6B**). We observed that T1L replicated faster than the clones, but there was a time dependent, modest increase of L1 expression in T1L SP B and T1L SP C. Together, these results suggest that T1L SP B and T1L SP C are less potent activators of the antiviral response and replicate more slowly than the parental T1L strain (**Figure 6B**). Although T1L SP B and T1L SP C elicited a weaker antiviral response than the parental T1L strain, this diminished innate signaling does not equate with reduced cytotoxicity. Instead, our data indicate that the clones are more cytotoxic in BCSCs despite their attenuated antiviral gene activation. This trend suggests that the parental T1L strain triggers a strong interferon response that may partially restrict its own cytotoxic potential. We posit that the SP clones may evade immune sensing and as a result, even though their early replication kinetics appear slower, the clones ultimately generate more cumulative cell stress and apoptosis at later time points. These findings highlight that strong antiviral signaling does not necessarily mean stronger oncolysis. Rather, reduced innate immune activation may facilitate more effective tumor cell killing. The schematic in **Figure 6C** summarizes the data gathered in this study. Although T1L infects BCSCs more efficiently, produces higher levels of viral protein, and triggers a stronger antiviral response 24 hours post infection, the SP clones ultimately induce more cell death. This enhanced cytotoxicity is accompanied by increased eATP, suggesting that the clones may promote a more immunogenic form of apoptosis. In addition, the T1L SP B generates more plaque-forming units than T1L 6 days post infection, indicating more effective viral spread and a potential bystander effect that could contribute to their greater killing capacity.

**Figure 6.**
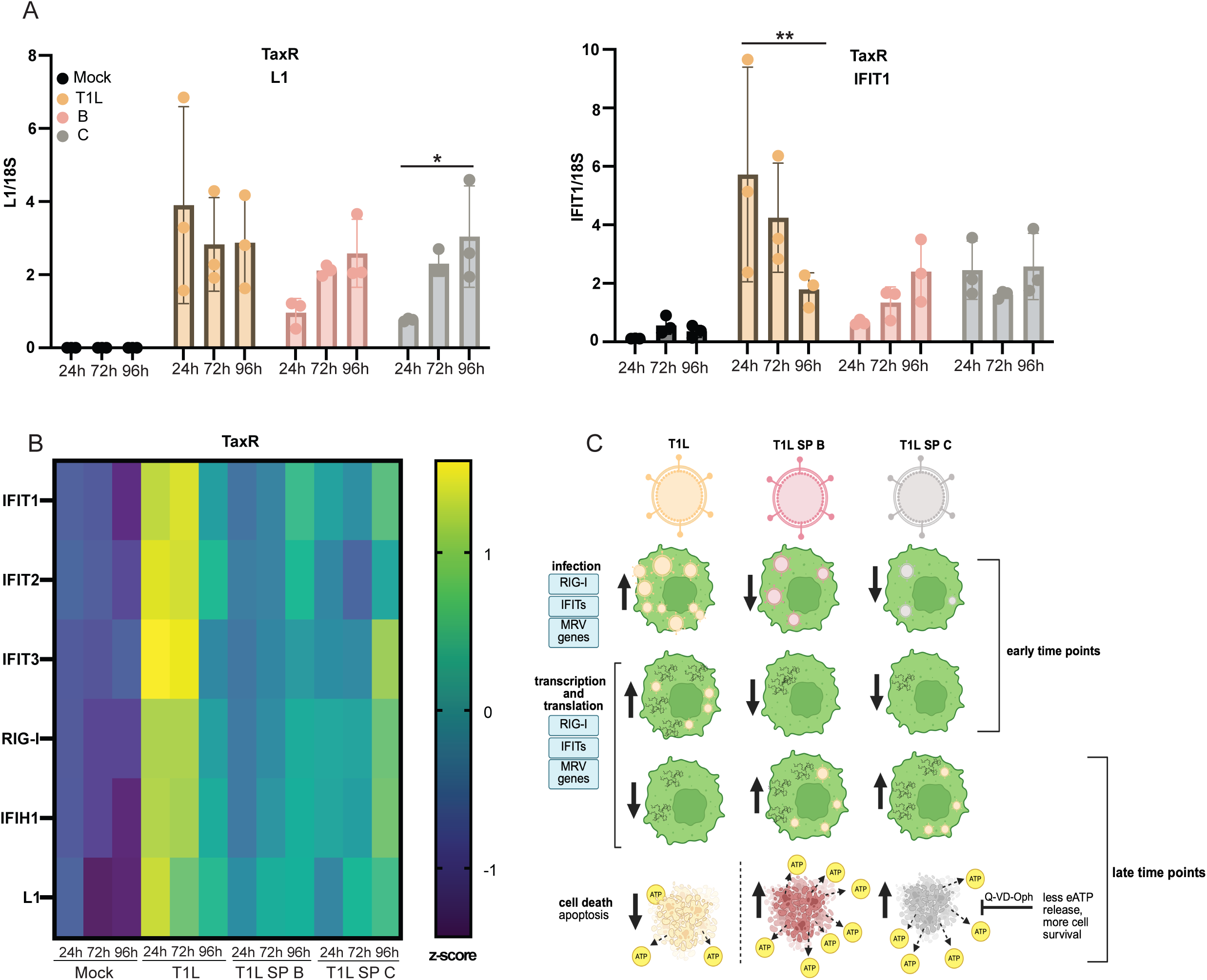
The viral clones elicit weaker antiviral responses compared to the parental T1L strain. **A**) qRT-PCR for L1 and IFIT1 on RNA isolated from MCF7 TaxR cells treated with MRV (T1L, SP B, and SP C)**. B)** Heatmap of qPCR results analyzed by normalization to 18S gene and subsequent z-score per gene n=3. **C)** Schematic of the proposed working model. Statistical significance is indicated as follows: *p ≤ 0.05, **p ≤ 0.01, ***p ≤ 0.001, and ****p ≤ 0.0001.

## Discussion

Our work explored the oncolytic effects of (3) MRV strains (T3D, T1L, R2) in ER+ breast cancer models under both 2D and 3D tumorsphere conditions, with the ultimate goal of identifying an MRV strain or variant that will target BCSC populations. In 2D conditions, all MRV strains reduced cell viability with no major differences between cell models. MRV Infection in MCF7 cells trended higher than the TaxR, but statistically significant changes were only observed with T3D (**Figure 1B**). In tumorsphere conditions, which enriches for BCSC populations, T3D, T1L and R2 elicited very modest reductions in tumorsphere viability (**Figure 1E**), highlighting the need for improving MRV virotherapeutics to better target BCSCs. We used serial passaging of T1L in a therapy-resistant, BCSC-enriched cell line (i.e. TaxR cells), to facilitate the identification of a BCSC-targeting MRV variant (**Figure 4A**). Importantly, this method produced several MRV clones with enhanced capacity to kill BCSC-enriched tumorspheres (**Figure 4C, D**).

### Transcriptomic Insights Into MRV-infected Cells

We used RNA sequencing of tumorspheres infected with T1L to provide insights into the effects of MRV infection in BCSCs (**Figure 2**), which has not been previously reported. Although only a few transcriptomic studies exist in response to MRV infection (57–63), some align with our work. For example, we detected pronounced antiviral gene expression, with elevated levels of interferon-stimulated genes (IFIT1, IFIT2, IFIT3), RNA degradation factors (OAS1, OAS3), and viral RNA sensors (DDX60, IFIH1, IFI44), among others (59,60). Moreover, we observed immune response pathways, such as type I interferon alpha (IFN-α) and type II interferon gamma (IFN-γ), significantly activated in MCF7 parental and TaxR BCSCs at 24h (**Figure 2D, 2E**), as well as the shared pathways (**Figure 2D**). Consistent with our findings, previous work examining oral MRV (RC402) in colon cancer tumor-bearing mice demonstrated upregulation of interferon-stimulated genes associated with both innate and adaptive immunity (62). In our data, the TaxR BCSCs unique pathway analysis suggests that both pro-viral (mitochondrial activity/RNA processing) and antiviral (Rho GTPase signaling) responses are likely occurring in parallel (**Supplementary S2 Figure E**). Consistent with this, Rho GTPase expression in host cells infected with viruses (e.g avian reovirus in Vero and DF-1 cells, measles virus in T cells, HIV in macrophages adenovirus in fibroblasts and others) has been shown to modulate viral entry, replication, and release, as well as innate immune sensing and inflammatory signaling (64–69). Upregulation of Rho GTPase signaling in TaxR BCSCs may also reflect MRV-driven cytoskeletal remodeling or a host response engaging in antiviral pathways. From our datasets of differentially expressed genes that are unique to MCF7 parental BCSCs or shared between MCF7 parental and TaxR BCSCs, we observed an upregulation of genes involved in innate immune signaling pathways and NF-κB activation through FADD RIP-1 pathway. The downregulated shared pathway analysis showed suppression of RNA processing, RNA modification, and nuclear transport pathways, which is consistent with MRV-induced host shutoff and cellular stress, whereby the virus suppresses mRNA maturation and nuclear-cytoplasmic trafficking to promote replication (70).

It should be noted that there are significant differences in basal gene expression when comparing MCF7 and TaxR BCSCs (**Figure 2E**). TaxR cells displayed a pre-activated antiviral immune transcriptional program at baseline, with strong interferon and inflammatory responses. A study in patient-derived pediatric brain tumorspheres found that baseline (uninfected) transcriptomic profiles could predict MRV sensitivity, as measured by EC₅₀ values, with sensitivity associated with mitochondrial activity and RNA processing, whereas resistance correlated with positive GTPase regulation, autophagy, and Ras protein signaling (71). However, in validation cultures they were not able to clearly separate the sensitive cells from the resistant ones (71). Our GSEA data suggest that high baseline interferon and inflammatory signaling in TaxR may underlie their resistance, an observation suggested by other studies (72–75). Notably, these immune-related pathways were not identified as predictive markers in Vazaios et al., 2024. Another study explored sensitivity and resistance to MRV (RV Jin-3) in uninfected pancreatic ductal adenocarcinoma (PDAC) patient-derived organoids (PDOs) where sensitivity was associated with proliferative, immune response, MAPK-activated, stem-cell-like transcriptional programs, and resistance was associated with granulocyte colony stimulating factor, mitochondrial processing, apoptosis and others (76). PDAC PDOs showed that MRV sensitivity was associated with certain immune-related programs alongside proliferative and stem-cell-like transcriptional states (76). In our system, TaxR cells displayed elevated baseline interferon and inflammatory signaling, which we hypothesize at least partially restricts viral replication.

### Adaptive Evolution by Serial Passaging of OVs

Our results support the hypothesis that serial passaging can enrich for viral variants with greater oncolytic potential than the parent virus. Other studies have employed viral adaptation by serial passaging with other viruses (77–79), including MRV (37,80–82). For example, propagation of T3D in JAM-A-negative glioblastoma cells developed mutants (RV Jin-1, Jin-2, Jin-3) arising from spontaneous mutations in the σ1 spike protein that allowed the virus to infect cells independently of JAM-A (82). In subsequent studies it was found that RV Jin-3 induces autophagy in mouse embryonic fibroblasts (83), exhibits antitumor effects, releases inflammatory cytokines, and induces immune stimulated genes in prostate cancer (84). RV Jin-1 has been shown to induce necroptosis in murine cells but not in human tumor cell lines (85). Building on the concept of virus adaptation, Rodríguez Stewart et al. developed a novel reassortant MRV, R2 (studied in **Figure 1**), by coinfecting TNBC cells with T1L, T2J, and T3D strains and using serial passaging to enhance viral adaptation (37). MRV-R2, which contains 9 segments of T1L and 1 segment (M2) of T3D, showed improved infectivity and cytotoxicity in TNBC cells compared to parental strains. The authors hypothesized that specific mutations in several viral genes (i.e. S4, that encodes for the viral factory protein σNS; S3, that encodes for the outer capsid protein σ3, and L3 that encodes for the inner capsid protein λ1) likely contributed to enhanced infection. MRV-R2 infection strongly induced type III interferon (IFN-λ), but not type I IFN responses, particularly in the presence of topotecan (37). A follow up study by Rodríguez Stewart et al., showed that R2 displayed superior infectivity and cytotoxicity in TNBC cells through suppression of MAPK/ERK signaling and induction of noncanonical, caspase-9–dependent cell death (86). Together, these observations suggest that adaptive changes in MRV can enhance cytotoxicity by increasing viral infectivity, modulating host immune responses, and triggering alternative cell death pathways, underscoring the utility of adaptive evolution for the development of more effective OVs.

Interestingly, in our study, the enhanced oncolytic activity of T1L SP B and T1L SP C was not due to enhanced infection efficiency as we observed that at 24h post infection fewer TaxR cells were infected. Our data suggests that the enhanced killing effect is likely due to an increase in viral production after primary infection. Over time, our plaque assays showed T1L SP B generates more viral particles (**Figure 4G**) and induces more cell death than T1L (**Figure 4D**). Because T1L SP B and T1L SP C exhibited enhanced cytotoxicity in TaxR tumorspheres, independent of infection rate or viral protein levels, we posit it is likely through apoptosis and bystander effects mediated by cytokines, death ligands, and DAMP release. T1L SP B in particular promotes increased eATP release (**Figure 5G**), indicative of immunogenic cell death, highlighting its potential as a selective BCSC oncolytic agent. Consistent with prior reports that MRV induces apoptosis in cancer cells via caspase activation (87) our results using pan-caspase inhibitors confirmed that T1L, T1L SP B, and T1L SP C rely on apoptosis to mediate cell death in both MCF7 and TaxR cells. T1L displayed a stronger antiviral response than the SP clones (**Figure 6A**), suggesting this strain is more easily detected by immune sensors and that the clones evoke a muted antiviral response. Further research is needed to determine the mechanism by which these clones elicit cytotoxicity.

### Oncolytic MRV in CSCs

Recent research highlights a connection between MRV susceptibility and stem-like cellular characteristics, suggesting that stemness may play a central role in determining viral efficacy. To our knowledge, Marcato et al, 2009 was the first published study which demonstrated that oncolytic MRV induces regression of human breast tumors in xenograft models and effectively targets BCSCs and non-BCSCs. Immunofluorescence and TUNEL assays confirmed that both BCSCs and non-BCSCs died via apoptosis (36). A recent study showed that MRV can infect and lyse cancer cells, CSCs, healthy embryonic stem cells (ESCs), and induced pluripotent stem cells (iPSCs), whereas adult stem cells (mesenchymal and hematopoietic) are largely resistant. Differentiated ESCs or iPSCs were resistant to MRV, demonstrating that the stem-like state promotes viral permissiveness. Furthermore, murine mammary cancer cells that were originally resistant to MRV infection became highly susceptible after reprogramming into iPSCs using Yamanaka factors, highlighting the link between stemness pathways and MRV sensitivity (88). Importantly, these studies used different strains (RV Jin-3, T3D) than those used in our work, which focused on T1L and the T1L SP clones. This distinction emphasizes that conclusions regarding stemness-mediated susceptibility and safety should be interpreted in the context of the specific viral strain and that results may not be directly transferable between MRV variants without further validation. Together, these findings emphasize the role of CSCs in modulating MRV susceptibility; however, further studies across additional MRV strains and different cancer models are needed to fully validate and extend these findings for the development of MRV-based OVs.

## Conclusions

This research shows that all three parent MRV strains (T3D, T1L, R2) infect and reduce cell viability in ER+ breast cancer models in 2D conditions, while in 3D tumorsphere models (enriched for BCSCs), infection and tumorsphere killing were lower. Through serial passaging we developed three MRV clones that replicate better in therapy-resistant tumorspheres and have superior oncolytic effects. This enhanced oncolytic effect is likely due to immune-mediated cell death, including the release of DAMPs. In summary, serial passaging of T1L MRV in BCSCs can create novel MRV variants with enhanced oncolytic activity. OVs hold significant promise in targeting BCSCs, offering a new approach to overcome tumor resistance and enhance long-term anti-cancer immunity.

## Declarations

### Ethics approval and consent to participate

Not applicable. This study used only established human and mouse cell lines and did not involve human participants, human tissue, or animal subjects.

### Consent for publication

Not applicable. This manuscript does not contain data from any individual person.

### Availability of Data and Materials

All data generated or analyzed during this study are included in this published article and its supplementary information files.

### Author’s Contributions

JHO, PD supervised the study. NIRO wrote the initial draft of the manuscript and conducted all experiments. PD performed the virus purification and plaque assays. NIRO, JH and EB contributed to the bioinformatics analysis. KM contributed to the analysis of immunofluorescence and western blotting quantification. All authors reviewed, edited, and approved the final manuscript.

## Supporting information

Supplementary Figures combined

## Acknowledgements

We acknowledge the support of the University of Minnesota (UMN) Flow Cytometry Resource, which allowed for the acquisition of experimental data. We thank Dr. Carol Sartorius for providing the UCD cell lines. We thank Dr. Bernardo Mainou for providing R2, motivating the initiation of these studies, and critical reading of the manuscript. We also thank Angela Spartz for critical reading and editing of the manuscript. We acknowledge Novogene, for conducting RNA sequencing and the Center for Genomics and Bioinformatics at Indiana University Bloomington for conducting viral RNA sequencing.

