## Supplementary figures and images for "Viral evolution of T1L mammalian orthoreovirus enhances breast cancer stem-like cell killing"

### Supplementary Figures combined

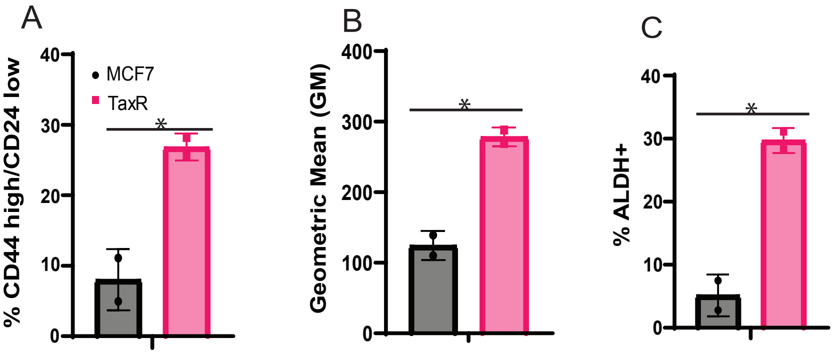

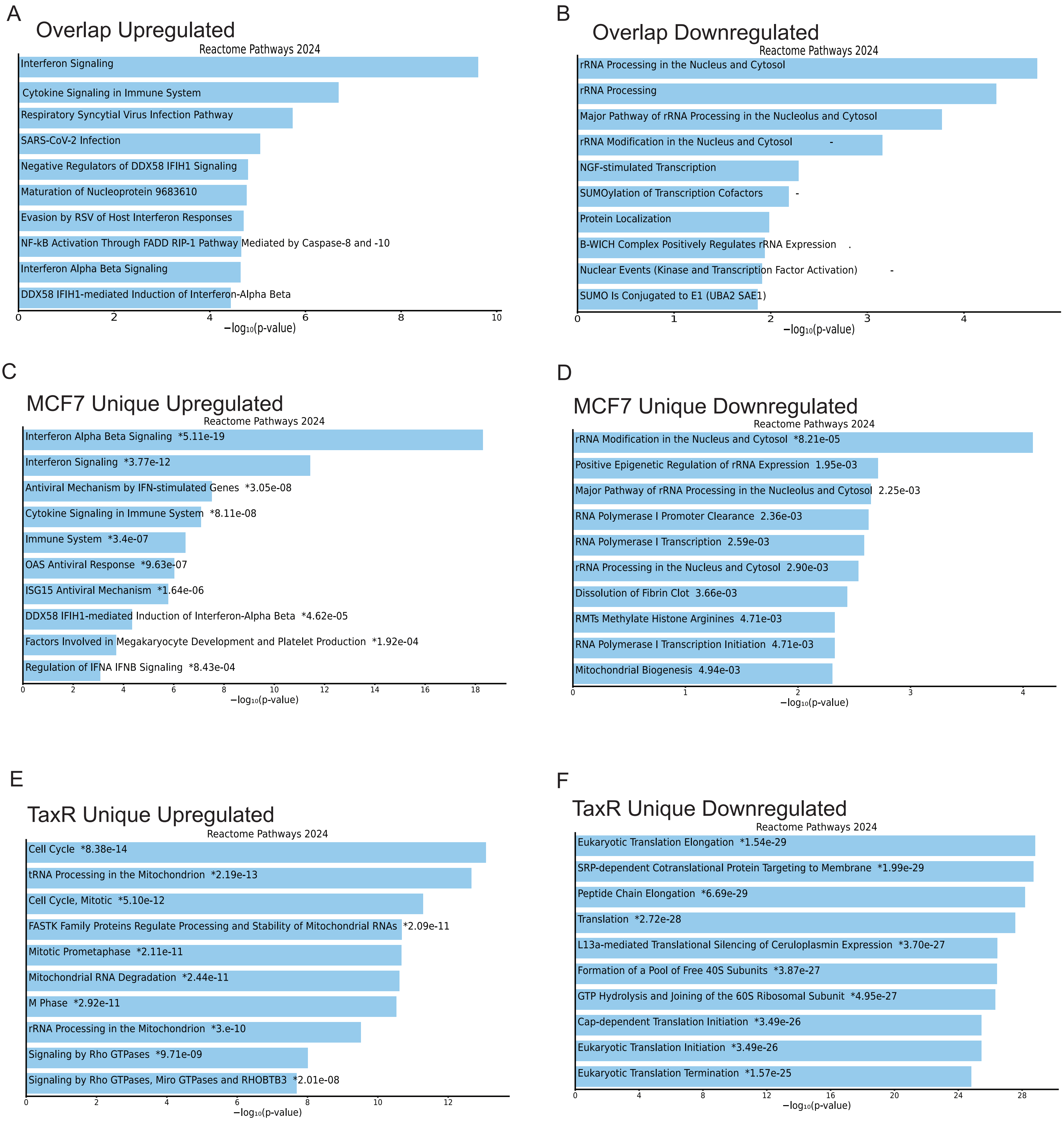

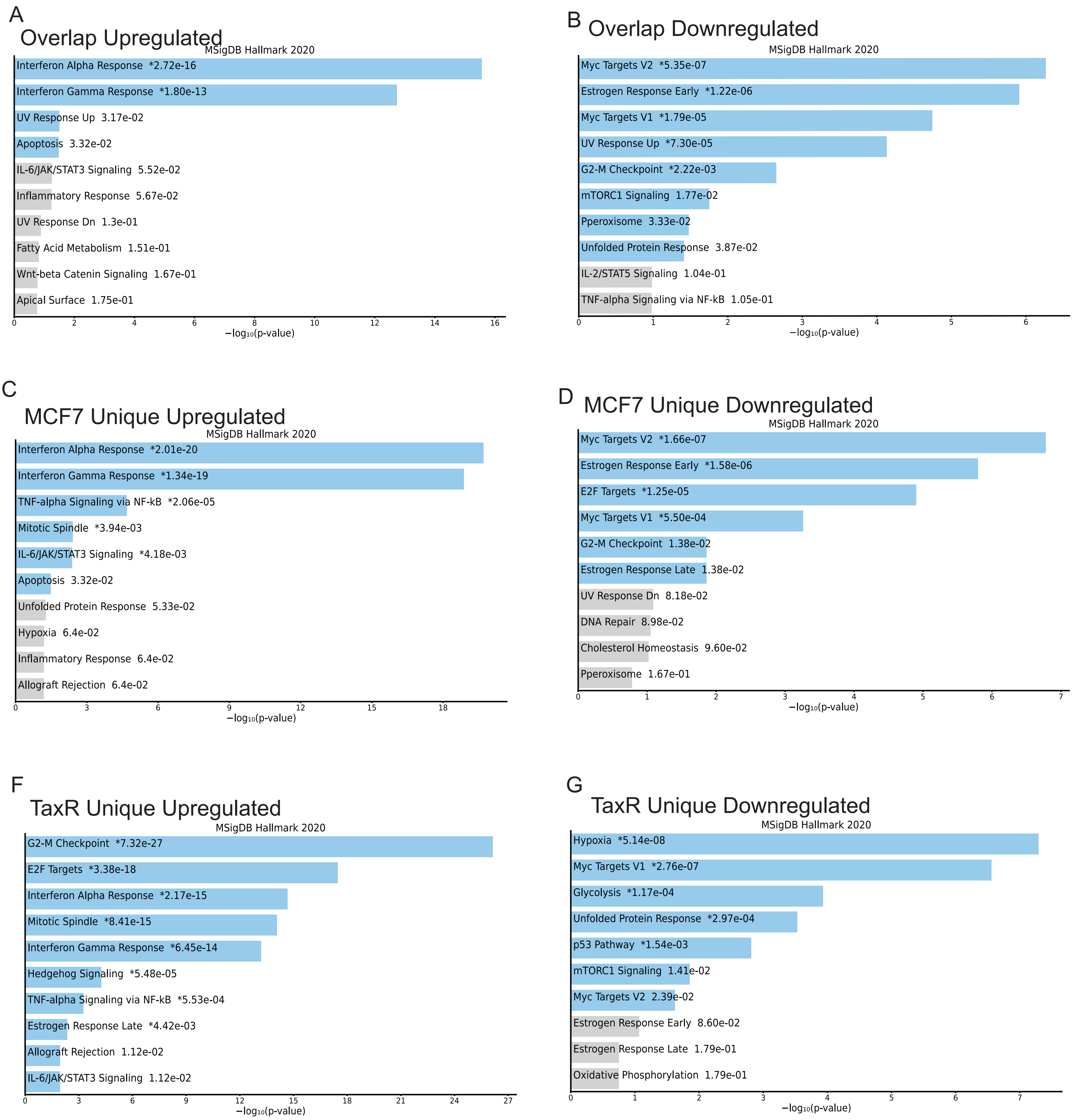

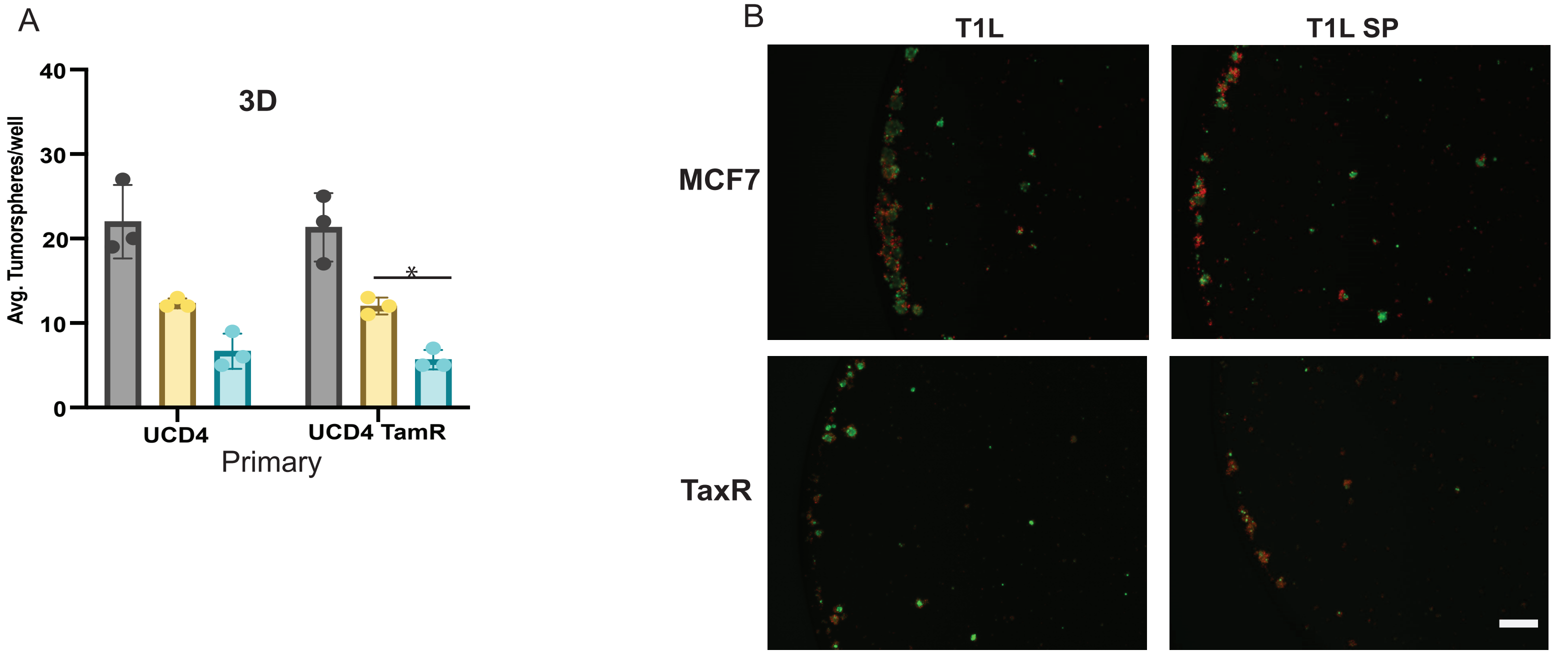

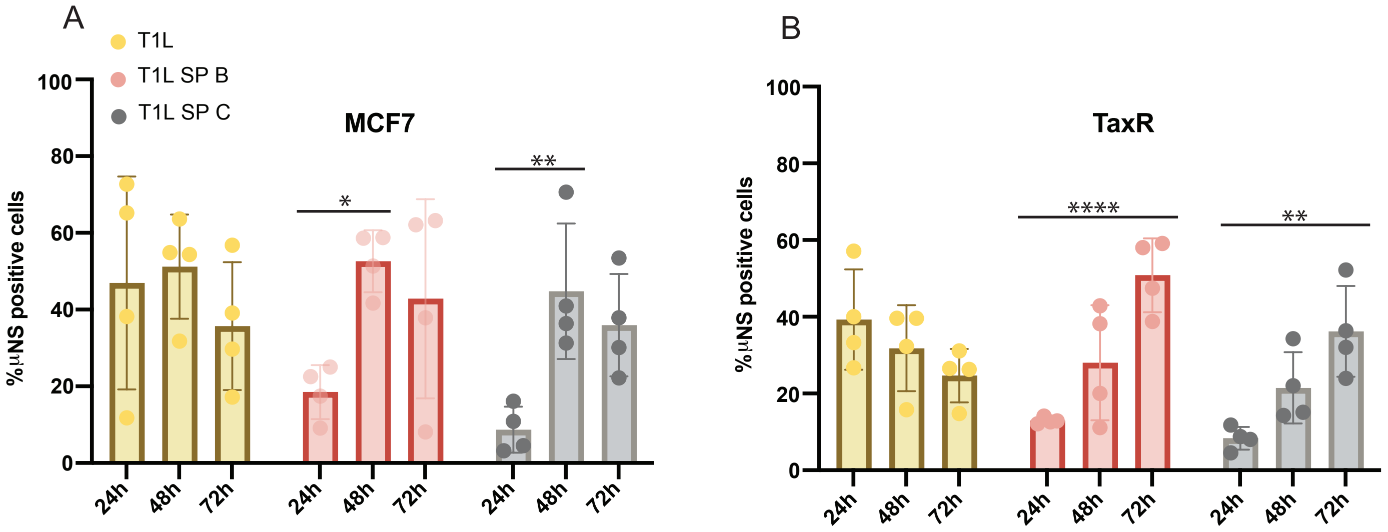

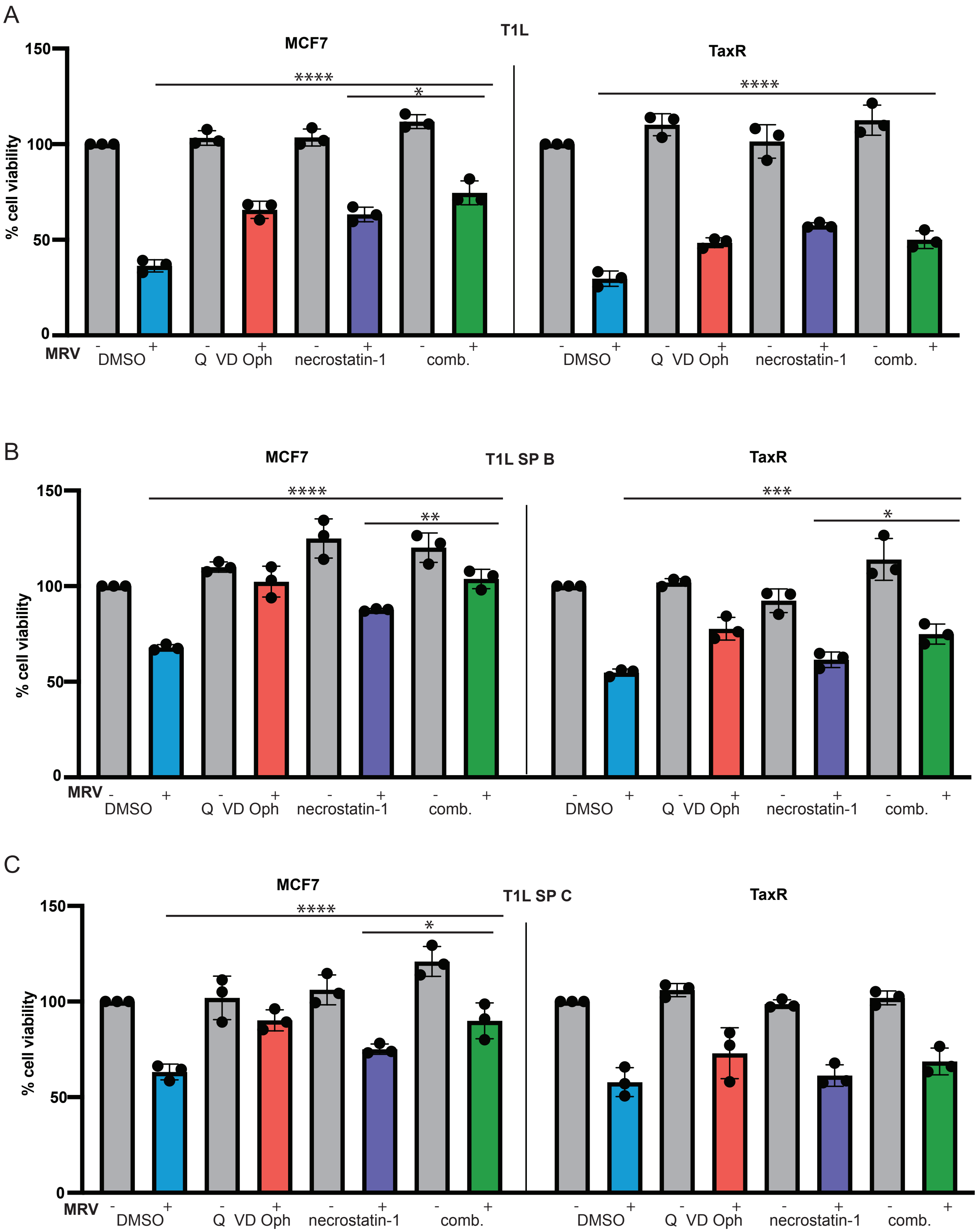
