## Supplementary Figure Legends for "Viral evolution of T1L mammalian orthoreovirus enhances breast cancer stem-like cell killing"

Supplemental Figure Legends

Supplementary Figure S1

**A)** Percentage of CD44^hi^/CD24^lo^ BCSCs using flow cytometry. Statistical Analysis: Unpaired T Test. **B)** Geometric Mean of ALDH activity by flow cytometry using the ALDEFLUOR kit. Statistical Analysis: Unpaired T Test. **C)** Percentage of ALDH activity in BCSCs by flow cytometry using the ALDEFLUOR kit. Statistical Analysis: Unpaired T Test. Statistical significance is indicated as follows: *p ≤ 0.05, **p ≤ 0.01, ***p ≤ 0.001, and ****p ≤ 0.0001.

Supplementary Figure S2

**A)** Reactome unique MCF7 upregulated pathways **B)** Reactome unique MCF7 downregulated pathways. **C)** Reactome unique TaxR upregulated pathways. **D)** Reactome unique TaxR downregulated pathways. Statistical significance is indicated as follows: *p ≤ 0.05, **p ≤ 0.01, ***p ≤ 0.001, and ****p ≤ 0.0001.

Supplementary Figure S3

**A)** Molecular Signatures Database (MSigDB) unique MCF7 upregulated pathways **B)** MSigDB unique MCF7 downregulated pathways. **C)** MSigDB unique TaxR upregulated pathways. **D)** MSigDB unique TaxR downregulated pathways. Statistical significance is indicated as follows: *p ≤ 0.05, **p ≤ 0.01, ***p ≤ 0.001, and ****p ≤ 0.0001.

Supplementary Figure S4

**A)** UCD4 and UCD4 TamR tumorspheres infected with mock, T1L or T1L SP, n=3, Statistical Analysis: Two-Way ANOVA. **B)** Representative images of MCF7 and TaxR tumorspheres infected with T1L and T1L SP. Statistical significance is indicated as follows: *p ≤ 0.05, **p ≤ 0.01, ***p ≤ 0.001, and ****p ≤ 0.0001.

Supplementary Figure S5

**(A–B)** MCF7 and TaxR cells were infected with T1L, T1L SP B or T1L SP C for 24h, 48h, and 72h. Viral factories were detected using a μNS specific antibody, and nuclei were counterstained with DAPI. Quantification was performed from four images per condition. Statistical analysis: Two-Way ANOVA. Analysis was conducted comparing each condition to the 24h time point. Statistical significance is indicated as follows: *p ≤ 0.05, **p ≤ 0.01, ***p ≤ 0.001, and ****p ≤ 0.0001.

Supplementary Figure S6

**(A–C)** alamarBlue assay in MCF7 and TaxR cells treated with DMSO, Q-VD-Oph (20 μM), necrostatin-1 (50 μM), or a combination of both inhibitors, in the presence or absence of viral infection. Cells were infected with T1L **(A)**, T1L SP B **(B)** or T1L SP C **(C)** n=3, Statistical analysis: Two-Way ANOVA. Statistical significance is indicated as follows: *p ≤ 0.05, **p ≤ 0.01, ***p ≤ 0.001, and ****p ≤ 0.0001.
