## Supplementary Table S1 for "Viral evolution of T1L mammalian orthoreovirus enhances breast cancer stem-like cell killing"

| Primer | Forward | Reverse |
| --- | --- | --- |
| IFIT1 | ATCTCAGTGAGGTCAGGTTTTCT | TGCATGCACACATTCAGTCC |
| IFIT2 | CCGAACAGCTGAGAATTGCAC | GCCCTCTTTGGGAACATAGC |
| IFIT3 | CAAAATCAACCGGGACCCCA | TCCTATCAGAGAAGCAGGGACT |
| IFIH1 | GACGTCTTGGATAAGTGCATGG | TCTGCAGCAGCAATCCGGT |
| RIG-I | CTAAGGGGATGATGGCAGGT | GGGCCAGTTTTCCTTGTCT |
| L1 | AACTGACAGATGCTCACCCG | CCGCTCGTCCAGATTTCGTA |

Supplementary Table 1. Primer sequences for qRT-PCR
